# A map of signaling responses in the human airway epithelium

**DOI:** 10.1101/2022.12.21.521460

**Authors:** Katherine B Mccauley, Kalki Kukreja, Aron B Jaffe, Allon M Klein

## Abstract

Receptor-mediated signaling plays a central role in tissue regeneration, and it is dysregulated in disease. Here, we build a signaling–response map for a model regenerative human tissue: the airway epithelium. We analyzed the effect of 17 receptor-mediated signaling pathways on organotypic cultures to determine changes in abundance and phenotype of all epithelial cell types. This map recapitulates the gamut of known airway epithelial signaling responses to these pathways. It defines convergent states induced by multiple ligands and diverse, ligand-specific responses in basal-cell and secretory-cell metaplasia. We show that loss of canonical differentiation induced by multiple pathways is associated with cell cycle arrest, but that arrest is not sufficient to block differentiation. Using the signaling-response map, we show that a TGFB1-mediated response underlies specific aberrant cells found in multiple lung diseases and identify interferon responses in COVID-19 patient samples. Thus, we offer a framework enabling systematic evaluation of tissue signaling responses.

## Introduction

The proper cell type composition of tissues is established through the action of extra-cellular signaling pathways, and changes in signaling occur ubiquitously in disease^1^. Establishing how different pathways modulate cell-type composition, organization and behavior therefore represents a priority in the fields of developmental biology and tissue physiology.

The question of how a tissue responds to extra-cellular signals is exemplified in the airway epithelium, a regenerative tissue exposed to continuous environmental stimuli yet demonstrating substantial long-term stability against perturbations^2,3^ The airway epithelium is composed of a basal stem cell pool that gives rise to six distinct mature cell types with roles in host defense and mucociliary clearance from the lung: club cells, mucin-rich goblet cells, multi-ciliated cells, pulmonary neuroendocrine cells (PNECs), tuft cells, and pulmonary ionocytes^2,4–6^. The relative abundance of each of these six cell types is actively regulated, and responds to diverse environmental insults: infectious and allergic stimuli lead to increased goblet cell numbers, while injury leads to differentiation of squamous cells at the expense of mucociliary cells^2,7^ and some cells undergo an epithelial-to-mesenchymal transition (EMT)^2,7^. Persistence of these cell states is associated with lung diseases^2^. It is a long-standing goal to identify signals that induce these changes in airway epithelial composition, and to better understand the effects of different ligands on the different epithelial cell types^2^.

Multiple extracellular signaling pathways modulate the composition of the airway epithelium including Notch, Wnt, transforming growth factor beta-1 (TGFB1), epidermal growth factor (EGF), bone morphogenetic protein 4 (BMP4), fibroblast growth factors (FGFs), and interleukins (IL13 and IL17)^2,7–12^ Other physiological cues, including mechanical strain^13,14^ and epithelial structure^15,16^ can also alter tissue composition. Several pathways have been shown to drive the primary modes of abnormal differentiation: persistent signaling through Notch, IL13, and IL17 induce goblet cell hyperplasia; persistent EGF signaling induces squamous metaplasia; and TGFB1 has been implicated in EMT and epithelial senescence associated with pulmonary fibrosis^2,3,12,17–21^.

Building on this extensive work, we set out here to construct a signaling–response map that could address questions that have until now been difficult to answer: (1) we still do not know how all but the best-studied pathways alter the abundance of all cell types, including rare cell types of the epithelium. The categorization of abundance changes into squamous- and secretory-cell metaplasia may lose information on how the epithelium re-organizes in response to different signals, and few signaling pathways have been evaluated for their potential to regulate the frequency of pulmonary ionocytes, tuft cells and PNECs despite their potential roles in disease^5,6,22–25^. (2) We do not know how each cell type changes in gene expression in response to different signals, and how plastic are the phenotypes of cells. Different ligands could induce similar (i.e. canalized or convergent^26^) phenotypic responses in a cell type, or not, while individual ligands could induce similar responses in different cell types, or act in a pleiotropic manner. And finally, (3) tissue atlases of human disease have recently identified disease-specific ‘aberrant’ cell states in tissues by single-cell RNA sequencing (scRNA-Seq)^27–30^ and it would be useful to relate these states to signaling pathways that may be potential targets for therapeutic intervention. A signaling– response map could address these three questions by defining changes in cell type abundance and gene expression in an unbiased manner, identifying changes not evident in the canonical gene repertoire used to study the tissue. We here make use of scRNA-Seq to construct such a version of such a map, encompassing responses to 17 signaling pathways.

To facilitate analysis of human disease atlases, it is desirable to study signaling responses in primary human cells. We made use of air-liquid interface (ALI) organotypic cultures of primary human bronchial epithelial cells (hBECs), which form a mucociliary epithelium containing physiological cell types. We evaluated changes in cell type composition in response to different signaling stimuli, and transcriptional responses to signaling. Through these analyses we identified stereotyped axes of variation in cell type composition, evaluated change in rare cell type abundance, defined convergent and unique transcriptional signatures of different pathways, and identified the cell types transcriptionally responding to different stimuli. We then explored the use of this signaling map to infer signal activation in lung diseases from patient tissue atlases. Thus, this study provides a framework for quantitative characterization of signaling responses, and it serves as a resource for predicting tissue-specific signaling signatures in diseases of the airway epithelium.

## Results

### Modeling human airway epithelial regeneration under stimulation of signaling pathways

To prioritize potential signaling pathways regulating the airway epithelium, we shortlisted receptors with cognate ligands^31^ and transcript expression enriched in hBECs relative to other human tissues (**Fig. 1A**). We did so by comparing RNA-Seq data from the GTEx Portal^32^ with scRNA-Seq data from human airway epithelial cells^5^, to identify 97 candidate receptors. From these we focused on those involved in immune, developmental, and hormonal signaling, and shortlisted a subset of 16 ligands and one chemical agonist known to interact with 31 of the 97 receptors (**Fig. 1A, S1A**,**B**). The selected ligands have previously been studied in varying contexts of airway epithelial differentiation and disease as summarized in **Table S1**.

**Figure 1.**
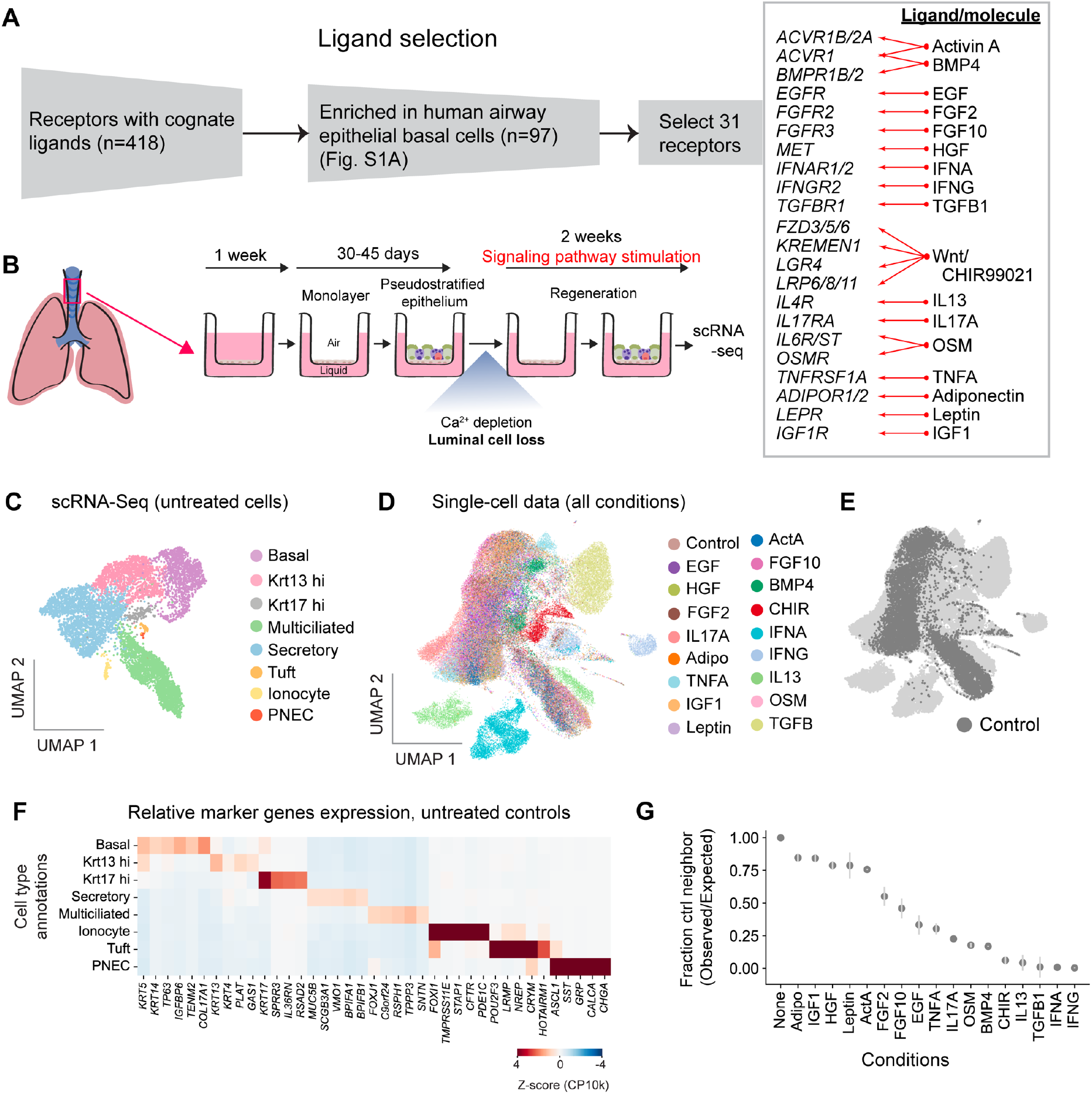
A single-cell map of receptor mediated signaling induction in human airway epithelial cells. **(A)** Approach for selection of signaling ligands for this study (for analysis details see **Fig. S1**). Red arrows show cognate ligand-receptor interactions. **(B)** Schematic of the organotypic regeneration assay for evaluating changes in hBEC differentiation under signaling stimulation. **(C**,**D**,**E)** UMAPs of scRNA-Seq data from **(C)** untreated cells, colored by cell type annotations, **(D)** all cells colored by treatment condition, and **(E)** all cells with control cells highlighted. **(F)** Expression of marker genes in annotated cell types in the control data quantified by scaled log_10_(CP10k + 1) where CP10k = Counts per 10k total counts. **(G)** De-mixing of treated cell states from untreated cell states quantified by the observed/expected ratio of nearest-neighbor cell fraction coming from control condition (100 nearest neighbors used). Lower values indicate separation of treated from untreated cells. Here and in all figures, CHIR = CHIR99021.

To test the effect of the selected signaling molecules on airway epithelial composition, hBECs from 3 human donors were treated as shown in **Fig. 1B**: the cells were first differentiated without treatment into a pseudo-stratified epithelium that recapitulates the physiological cell types of the airway^5^, after which the luminal cell were stripped through calcium depletion, leaving the remaining basal cells to regenerate the tissue. This was done in the presence of a signaling agonist added to each well at saturating dosage (concentrations in **Fig. S1B**). For 16 pathways we applied purified ligands, and for one – the canonical Wnt pathway – we applied a small molecule agonist, a GSK3 inhibitor (CHIR99021) (**Fig. 1A**). After 2 weeks of differentiation, the final composition of the tissue was analyzed by scRNA-Seq and imaging.

As a technical control for the efficacy of the ligands, we tested whether the expression of induced transcriptional targets (1-3 per pathway, **Table S2**) was significantly increased (**Fig. S1C)**. For 14 out of 17 pathways, induced transcriptional targets increased after stimulation (family-wise error rate<0.05). Previously reported targets of three treatments (ActA, FGF2 and IGF1) did not change with statistical significance after multiple hypothesis correction, but they still showed an increase in average expression. We cannot rule out the possibility that these ligands did not activate their respective pathways.

### Signaling responses extend the transcriptional landscape of the airway epithelium

After filtering for low quality cells, we obtained 77,568 cell transcriptomes over the 18 conditions (17 treatments, and one control) represented by one donor (IFNG, OSM), two donors (ActA, Leptin) or three donors (control, and remaining 13 treatments). To obtain a first view of the data, we performed batch correction between donors^33,34^, and then generated UMAP embeddings for the control data (**Fig. 1C)** and for the full data set (**Fig. 1D,E**). In untreated controls, we observed clusters representing all major airway cell types (**Fig. 1C**), indicating that the ALI cultures after luminal stripping fully regenerate the mucociliary epithelium. The clusters expressed canonical markers as expected (**Fig. 1F**) – *KRT5*+ basal cells, *MUC5B*+ secretory cells, *FOXJ1*+ multiciliated cells, *FOXI1*+/*CFTR*+ ionocytes, *ASCL1*+/*SST+* PNECs and *NREP*+ tuft cells. The untreated cells also defined two additional basal-like states: a *KRT13*+ state that expressed intermediate levels of basal and luminal keratins (KRT5 and KRT8 respectively) and variably expressed *KRT4*; and a separate, rare group of cells (138/7564) expressing low levels of both basal and luminal keratins (KRT5/8) and enriched for *KRT17* (**Fig. 1C**). The *KRT13*+ state does not relate to any classical airway epithelial state but appears transcriptionally transitional (basoluminal) and is homologous to a cell state found in mouse and human primary tissue samples^5,6,35^. The *KRT17+* cell state did not resemble any cell state of the mucociliary epithelium sampled from healthy mice or humans^5,6,35^, but *KRT17*+ cells have recently been reported in bronchial samples from patients diagnosed with pulmonary fibrosis^27,30^ and we have previously identified these states in HBEC ALI cultures^5^. In total, the transcriptomes of the cell types emerging in the regenerating cultures, and their abundances, establish that the regeneration assay recapitulates physiological cell types and sets a baseline against which to interpret responses to signaling pathway stimulation.

The UMAP visualization of the full data set (**Fig. 1D-E, S1D**) offers a simplified first view of the many changes in response to the 17 signaling ligands. These plots suggest that several treatments (ActA, Leptin, HGF, IGF1, and Adiponectin) gave rise to cells that were transcriptionally similar to untreated cells, while others (IFNA, IFNG and TGFB1) gave rise to cells that were not. However, the usage of UMAPs to construct 2D embeddings may lead to the loss of axes of variation in complex tissues that are heterogeneous in differentiation state, cell cycle state, and signaling response. Hence, we looked at the differences between signaling responses by calculating the average density of untreated cell transcriptomes in high-dimensional gene expression space (**Fig. 1G**). This analysis confirmed and quantified the trends seen in the UMAP plots: the smallest and largest deviations from untreated states were respectively associated with those treatments showing the least and greatest separation from untreated states on the UMAP. In what follows, we dissect these changes as they manifest (1) in cell type abundance, and (2) in altering transcription in response to signaling pathway stimulation.

### Mapping changes in cell type abundance upon signal pathway stimulation

To evaluate how pathway stimulation altered the abundance of epithelial cell types, we assigned the cell transcriptomes from treated conditions into annotated cell types using a classifier trained on the untreated cells (**Fig. 2A**). This classification was refined by k-nearest-neighbor voting, and a final filtering step based on a requirement that cells classified to each cell type show enriched expression for associated marker genes (**Figs. 2B, S2A**). With these annotations, we calculated the frequency of cell types in each condition (**Fig. 2C**).

**Figure 2.**
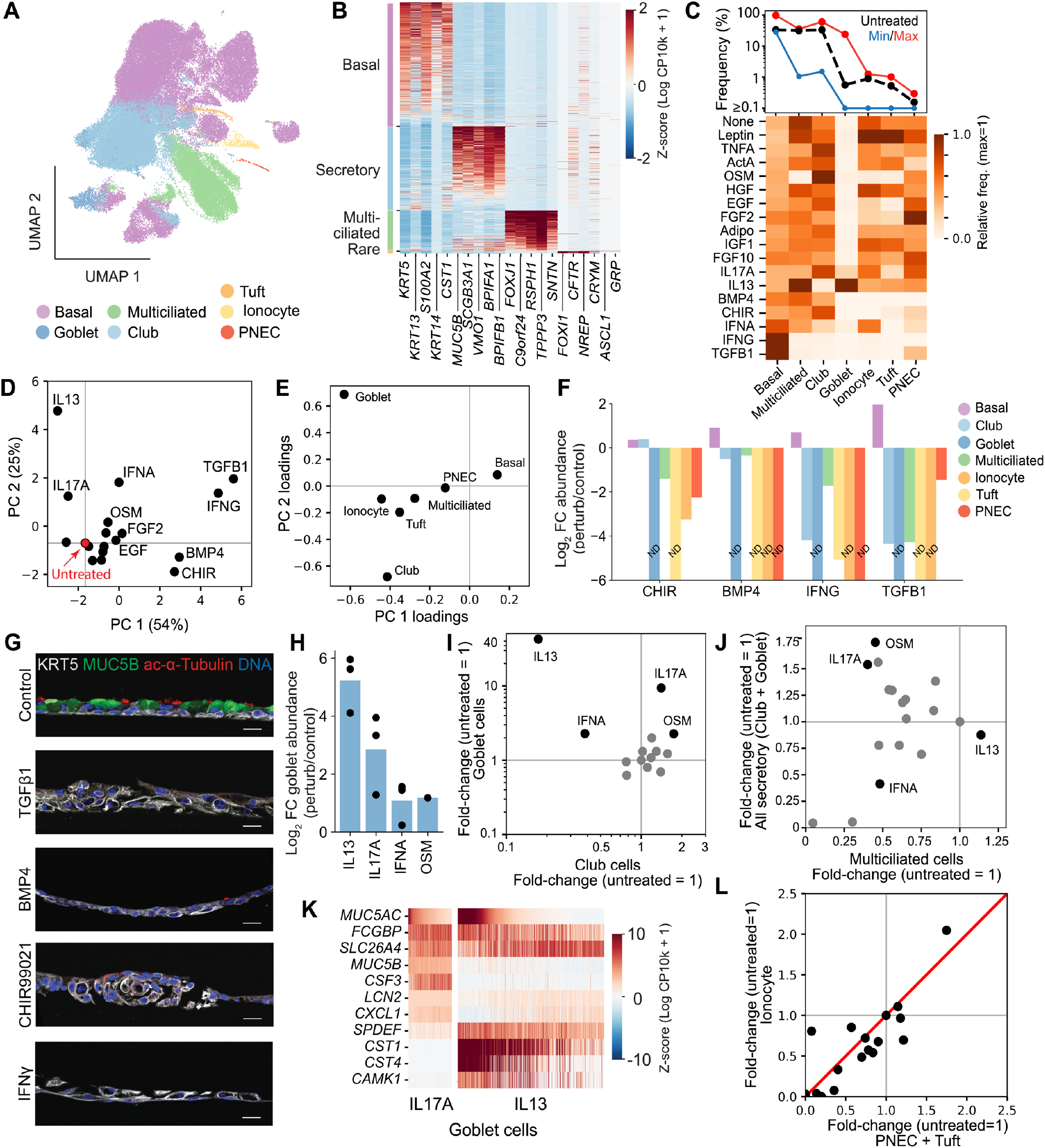
Distinct modes of basal and secretory cell metaplasia in response to different signals. **(A)** UMAP of all scRNA-Seq data from **Fig. 1D**, colored by canonical cell-type annotations learnt from untreated controls. **(B)** Gene expression heatmap showing that the classified cells across treatments preserve expression of marker genes for their respective cell types. Each row represents a single meta-cell showing average expression of 10 nearest neighbors; classified cell types on left. **(C)** Frequency of cell types after perturbation. *Top:* dynamic range of signaling-induced changes. Red=maximum; Blue=minimum; Black=untreated baseline. *Bottom:* heatmap of donor-averaged cell type frequencies in all conditions. **(D,E)** First two Principal Components of the cell type frequency matrix, after per-donor normalization, showing **(D)** values for each treatment, and **(E)** cell type loadings. PC1 corresponds to basal cell metaplasia, and PC2 corresponds to goblet cell hyperplasia. **(F)** Fold change in cell type frequencies for four conditions with highest PC1 values corresponding to loss of canonical differentiation. ND = Not detected **(G)** Representative immunofluorescence images of cross-sections of differentiated HBEC cultures treated with indicated cytokines and stained for KRT5 (white), MUC5B (green), and acetylated alpha-tubulin (red). Scale bars, 25 µm. **(H)** Fold change goblet cell abundance for four conditions with highest PC2 values, all showing >2-fold increase in goblet cell frequency. Points = donors; bar = mean. **(I)** Comparison of changes in frequency of goblet and club cells. Conditions inducing goblet cell hyperplasia are highlighted (black); remaining conditions shown in gray. See also **Fig. S2C. (J)** Comparison of changes in frequency of all secretory cells and multiciliated cells in the context of goblet cell hyperplasia. Colors as in (I). See also **Fig. S2D. (K)** Gene expression heatmap showing differences in goblet cell states induced by IL13 and IL17. Each column is a single meta-cell as in panel B. CP10k = Counts per total 10k counts. **(L)** Comparison of changes in frequency of ionocytes and PNEC+Tuft cells across all conditions, indicating tandem variation in the frequency of rare cell types.

The cell type frequencies in **Fig. 2C** vary across multiple treatments as compared to their control values, and when considered independently multiple changes are found to be statistically significant (85 null hypotheses rejected by Fisher’s Exact test at 5% FDR, donors p-values integrated by Fisher’s method; **Fig. S2B, Table S3**). To identify patterns in these many changes, we carried out a principal component (PC) analysis of the normalized cell type frequency matrix (**Fig. 2C, Table S3**). The first two PCs account for 79% of donor-normalized, log-fold-change variation in cell type frequencies (**Fig. 2D**) and they spontaneously recapitulate two well-known axes of airway epithelial metaplasia: PC 1 (54% of the variation) corresponds to expansion of basal-like cells at the expense of luminal cells, and PC2 (25% of the variation) describes goblet cell hyperplasia (**Fig. 2E**). The full observations, however, reveal differences within each of these two conditions, as we describe here.

PC 1 shows an expansion of basal cells and associated suppression of normal luminal differentiation with increasing severity, CHIR → BMP4 → IFNG → TGFB1 (**Fig. 2D**) consistent with prior studies carried out individually for each of these pathways (**Table S1**), but with differences in the degree of loss of each cell type (**Fig. 2F**). CHIR-treated cells permitted club cell differentiation but repressed multiciliated and goblet cells; BMP4 led to a modest suppression of mucociliary cells; while TGFB1 and IFNG led to a near-total loss of all differentiated luminal cells. Whether these pathways only suppress the differentiation of secretory and multiciliated cells, or also lead to loss of rare cell types (ionocyte, PNEC and tuft cells), was until now not known. We found that all the above-mentioned conditions led to depletion of rare luminal cell types, and with CHIR and BMP4 depleting the rare cell types to a much higher extent than secretory and multiciliated cells [6% (CHIR) and 30% (BMP4) reduction in total mucociliary cell fraction, compared to 91% (CHIR) and 100% (BMP4) loss of total ionocytes, PNECs and tuft cell fraction] immunostaining of ALI cultures for luminal and basal markers for IFNG, TGFB1, CHIR and BMP4 (**Fig. 2G**).

PC 2 identified several conditions leading to increased goblet cell abundance. As expected, the largest increase in goblet cell frequency was observed following IL13 treatment (36-fold expansion; FDR < 0.001), and we also observed expansion in response to IL17A^12^, OSM^36^ and IFNA^12^ (**Fig. 2H**). The effect of IFNA is consistent with prior work suggesting it can stimulate secretory cell differentiation^12^, and contrast with other reports showing reduced secretory differentiation in murine airway cells exposed to IFNA^37^. It is possible that inconsistencies in prior work could be explained by differences in how goblet cells expanded: IL13-mediated goblet cell expansion occurred at the expense of club cells (**Fig. 2I**), but not multiciliated cells (**Fig. 2J**). By contrast, IL17A increased both club and goblet cell frequency while the frequency of multiciliated cells was reduced, while IFNA increased goblet cell frequency while both club and multiciliated cell numbers were reduced (**Fig. 2I,J**). Thus, these ligands may act at different stages of mucociliary differentiation, and their effect may be missed by staining for pan-secretory markers. The goblet cells produced in each of these conditions were transcriptionally distinct and differed in expression of canonical goblet cell markers *MUC5AC, SPDEF* and *MUC5B*, as well as other genes (**Fig. 2K**). Together, these results suggest that goblet cell hyperplasia is not a monolithic phenotype: it encompasses multiple states of tissue composition and secretory cell phenotypes.

We also examined changes in the frequency of ionocytes, tuft cells, and PNECs across treatments. Multiple conditions led to a loss of these rare cell types (**Figs. 2C, F**) but it was striking that none of the conditions we examined led to statistically significant increases in any of these cell types (**Fig. S2B**). Changes in the rare cell types could be hard to evaluate because of their low frequency (∼1%, **Fig. 2L**), which reduces statistical power in identifying changes in their abundance in this study; however, we noticed that the ratio of ionocyte to tuft and PNECs remained roughly uniform across conditions (**Fig. 2L**, Pearson correlation R=0.85), consistent with their frequencies not being modulated independently of each other by any of the pathways studied here.

In summary, the analysis of canonical cell type abundances supports that (1) activation of several signaling pathways leads to loss of normal differentiation and expansion of basal-like cells with signaling-specific differences in the loss of different luminal cell types; that (2) goblet cells expand in several conditions that induce major differences in goblet cell gene expression, as well as differences in club and multiciliated cell frequencies; and that (3) none of the stimuli studied expanded rare cell types.

### Convergent and unique transcriptional responses to signaling

We next sought to evaluate the transcriptional response to each signaling stimulus. Discovery of differentially expressed genes (DEGs) between treated and untreated cells for each cell type (rank-sum testing, 5% FDR; **Table S4**) revealed that the cell state consistently showing the largest changes after stimulation were those classified as basal (**Fig. 3A**). This bias in transcriptional response across all treatments may reflect the plasticity of undifferentiated basal cells to undergo alternative modes of differentiation. Among all conditions, the largest responses were seen upon treatment with IFNG, IFNA, IL13, CHIR and TGFB1 (> 100 genes, FDR < 0.05). IFNA stood out as inducing a response in all cell types, whereas five of the 17 ligands (ActA, Adiponectin, IGF1, HGF, FGF10) induced few differentially-expressed genes in any cell type.

**Figure 3.**
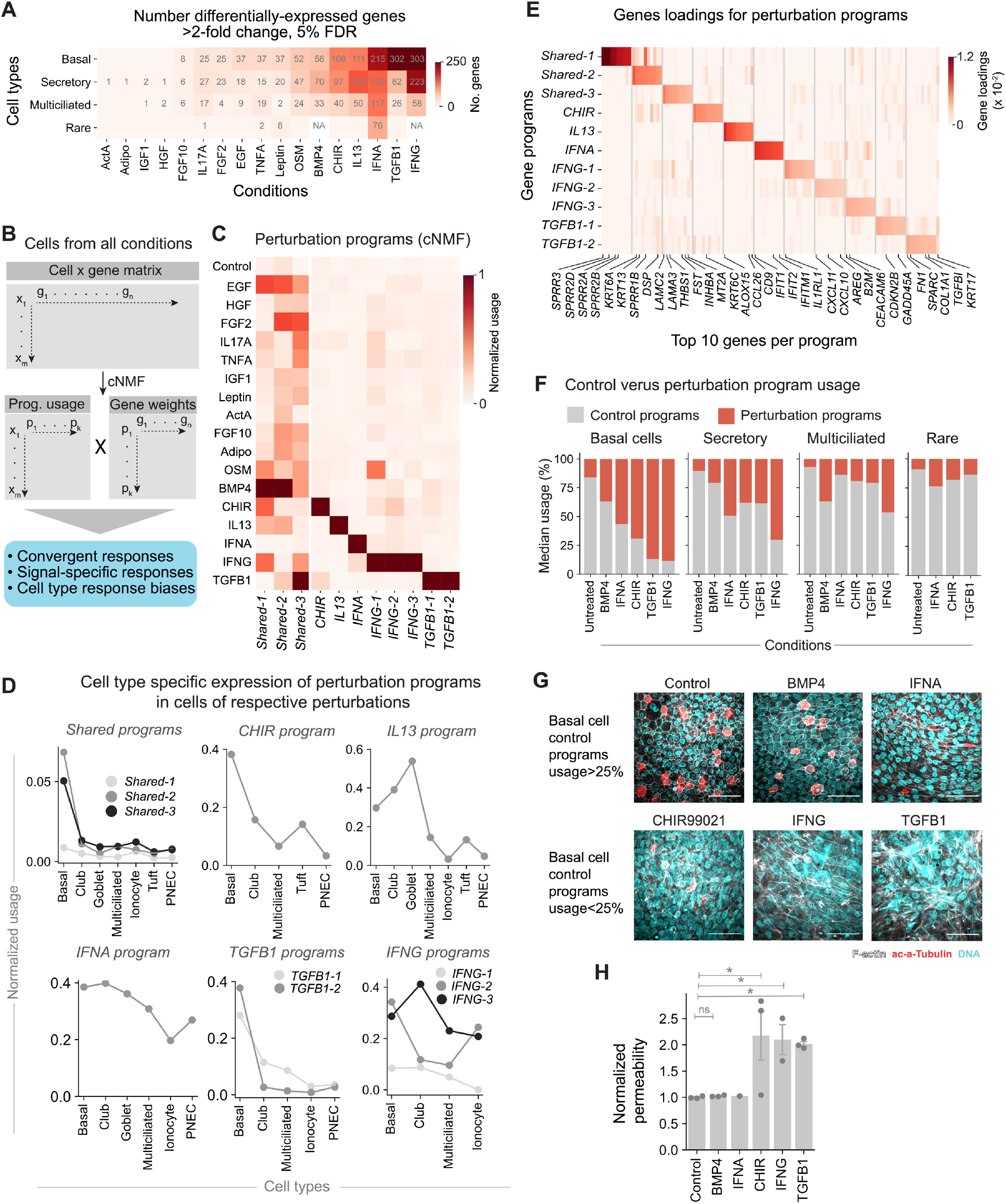
Evidence of convergent and pathway-specific transcriptional responses including loss of canonical cell identity. **(A)** Number of genes showing >2-fold differential expression in basal, secretory (club, goblet), multiciliated and rare (ionocyte, tuft, PNEC) cells following each treatment. Empty boxes = no genes; N/A = no cells present. **(B)** Schematic for gene program analysis of signaling responses. **(C)** Mean usage of eleven treatment-induced gene programs across all cells from each treatment condition the presence of convergent (*Shared-1-3*) and pathway-specific programs (remaining programs). A further nine control programs are shown in **Fig. S5A**. The heatmap is first column-normalized (sum=1) and then row-normalized (max= 1). **(D)** Transcriptional responses vary across cell types as seen from the mean usage of signaling programs. For shared programs, usage is averaged across all treatments; for perturbation-specific programs, usage is calculated for cells from one perturbation. **(E)** Loadings of top 10 genes for each of the signaling induced program (left) and of epithelial KRT genes (right) across all signaling programs; full table for gene loadings is provided in **Table S5. (F)** Putative loss of canonical basal cell identity, but not mucociliary cell identities, is observed by the near-complete replacement of control transcriptomic programs (grey) in basal cells by induced programs (red) in response to CHIR, IFNG, and TGFB1. Other treatments are shown for contrast. **(G)** Representative immunofluorescence images of whole-mount differentiated HBEC cultures treated with indicated cytokines and stained for F-actin (white), acetylated-alpha-tubulin (red), and DNA (Hoechst; blue). Scale bars, 50 µm. Bottom row conditions, corresponding to loss of basal cell identity in panel F, show cytoskeletal disorganization. **(H)** Quantification of epithelial permeability measured by transit of lucifer yellow dye across the epithelial surface of cultures treated with indicated cytokines. Bars represent mean ± SEM, n = 3 HBEC donors (IFNG: n=2; IFNA: n=1) biological replicates shown as points, normalized to untreated fluorescence = 1. * = p-value ≤0.05 by Wilcoxon rank sum test.

To gain insight into the nature of these transcriptional responses, we factorized the full scRNA-Seq data matrix into programs with variable usage across cells by consensus non-negative matrix factorization (cNMF)^38^ (**Fig. 3B**). Unlike DEG identification, matrix factorization is agnostic to both perturbations and cell types and so it offers a tool to define gene programs that recur across multiple cell types or stimuli, without the typical loss of sensitivity seen in DEG analysis for rare cell types. We defined 20 cNMF programs from our dataset, of which 9 were associated with unperturbed mucociliary epithelium (**Fig. S3A-C**) and 11 programs induced by signaling perturbations (**Figs. 3C-E, Table S5**). We do not discuss the control programs further, and instead focus on the transcriptional programs induced by the signaling perturbations.

The 11 perturbation programs collectively describe transcriptional changes in response to all signaling stimuli. Their usage pattern across conditions reveals a logic that is simple: of these programs, three were induced by multiple perturbations (**Fig. 3C**, *shared programs 1-3*). The remaining programs appeared only in response to one or two signaling conditions. We named these programs by the signaling condition that induces them (**Fig. 3C**).

The cNMF analysis also defines which cell types induced each of the different response programs (**Fig. 3D, S3C**), and the genes that define them (**Fig. 3E, Table S6**). The three shared programs were most enriched in basal cells (**Fig. 3D**). One of these programs (*shared-3*) was defined by increased expression of laminin genes, metallotheinin (MT2A), and inhibitors of TGF family members (Follistatin and Inhibin A), potentially indicating changes in extracellular matrix (ECM) properties and signaling (**Fig. 3E**). The two other convergent programs (*shared-1 and shared-2*) were enriched in genes observed in squamous epithelial tissues and which have been used as markers of squamous metaplasia in the airway: the Small Proline Rich Protein (*SPRR*) family in *shared-1*, and *SPRR, DSP* and *KRT6A* in *shared-2* (**Fig. 3E, Table S6**). These shared programs may thus represent distinct forms or progressive stages of basal cell differentiation into a squamous-like epithelium, and indeed they were induced by ligands that led to loss of luminal cell types and emergence of squamous-like morphologies (BMP4, IFNG, TGFB1) (**Fig. 2G, 3C**).

For the ligand-specific programs, the response across cell types was more varied (**Fig. 3D**): the two programs induced by TGFB1 (*TGFB1-1, TGFB1-2*), the program induced by CHIR (*CHIR*), and one of the programs induced by IFNG (*IFNG-2*) were most strongly induced in basal cells. However, the IFNG programs were also induced in the luminal cells still present after IFNG treatment, with one (*IFNG-3*) showing maximal expression in secretory cells. For IL13, the transcriptional response was maximal in goblet cells but the same program was also induced in club cells, multiciliated cells and basal cells. Thus, the IL13 transcriptional response is not directly a measurement of increased goblet cell numbers; it includes a large number of genes including the lipoxygenase *ALOX15*, whose activity promotes goblet cell differentiation in human airway^39^. For IFNA, the specific response was induced across all cell types, as expected from the DEG analysis (**Fig. 3A**). Of the genes upregulated by these programs (**Fig. 3E, Table S6**), we highlight that program *TGFB1-2* identifies cells that induced canonical markers of epithelial-to-mesenchymal transition (*SPARC, CDH2, FN1*), whereas *TGFB1-1* did not and corresponds to induction of diverse genes including cell cycle inhibitors (*CDNK2B, GADD45A*). Thus, these programs clarify the complexity in the responses to the different ligands, and they decouple convergent squamous-like responses from TGFB1-induced EMT-associated phenotypes and other pathway-specific phenotypes. The programs also clarify the pattern of expression of common markers used in tissue staining. Both TGFB1 programs, for example, induced expression of the cytokeratin *KRT17* (**Fig. 3E**), but the map revealed *KRT17* to also be induced by several other programs including the convergent program *shared-2* (**Fig. 3E, Table S6**).

### Loss of basal cell identity correlates with loss of epithelial barrier integrity

A question that can be asked from a systematic analysis of signaling responses is whether some ligands induce programs that qualitatively change cell identity. Definitions of canonical cell types historically depended on cell and tissue morphology, and on expression of marker genes including lineage-specifying transcription factors or unique structural proteins such as keratins. With access to whole-transcription information, we wondered whether the extent of remodeling of the cell transcriptome could offer an alternative and unbiased way of defining departures from canonical cell types. To formalize this idea, one can examine the expression of ‘control’ programs – those that specify the transcriptional state of cells in absence of treatment. Without treatment, control cNMF programs (defined in **Fig. S3A** and **Table S5**) composed >90% of the median transcriptome of luminal cells, and >80% of the median basal cell (**Fig. 3F**), while after treatment, control programs in basal cells exposed to TGFB1 and IFNG was almost entirely lost (median usage 13% and 11% respectively). BMP4, IFNA also led to somewhat reduced basal cell control program usage, but to a lesser extent, and CHIR represented an intermediate case. The residual luminal cells in all conditions continued to express control programs (>50% median usage in multiple conditions) (**Fig. 3F**) suggesting that luminal cells tend towards a more canalized identity in the face of perturbations as compared to basal cells. The near-complete loss of untreated transcriptional programs in basal cells suggests that TGFB1 and IFNG lead to a qualitative change in cell identity.

We expected that these large changes in whole-transcriptome state in response to TGFB1 and IFNG, together with the loss of luminal cells in these conditions, might be readily evident in the morphology and epithelial barrier function of the tissue after treatment with these ligands, but not after BMP4 or IFNA treatment. Staining the treated tissues for F-actin indeed revealed disorganization of the epithelial tissue in response to IFNG and TGFB1 (**Fig. 3G**). Further, dye-transport assays revealed a loss of epithelial integrity in response to these two treatments, while BMP4- and IFNA-treated cells had intact barrier function (**Fig. 3H**). As tight junctions form at the apical surface of polarized cells, this is consistent with BMP4/IFNA permitting differentiation of sufficient polarized luminal cells^40,41^. CHIR showed loss of epithelial integrity comparable to that seen for IFNG and TGFB1 (**Fig. 3G**). The loss of basal cell identity as seen from whole-transcriptome analysis together with alterations in luminal cell abundance thus correspond to changes in epithelial cell organization.

### Loss of differentiation induced by multiple pathways is accompanied by cell cycle arrest

Commonalities in the responses to perturbations can be used to identify shared mechanisms of action^42^. Here, we noticed that the conditions that led to loss of luminal differentiation (TGFB1, IFNG, BMP4 and CHIR) showed a downregulation in a control cNMF program associated with cell cycle (**Fig. S3A, 4A**), and an upregulation of CDK inhibitors (**Fig. 4B**). One hypothesis is therefore that cell cycle arrest is associated with loss of mucociliary differentiation. Indeed, it has previously been observed that cell proliferation in airway epithelial cells is suppressed by BMP4^43^, TGFB1^44,45^, and IFNG^37^. We tested here whether the decrease in cell cycle gene expression and increase in CDK inhibitor expression during regeneration after luminal stripping was indeed concurrent with cell cycle arrest by assaying cell number and EdU incorporation in these four conditions. Within 48 hours of epithelial stripping, we observed decreased epithelial cell density and loss of EdU incorporation **(Figs. 4C-E)** upon treatment with TGFB1, IFNG, BMP4 but not CHIR. Given this association between loss of normal differentiation and cell cycle arrest in three out of four conditions, we next asked whether cell cycle arrest is sufficient to alter differentiation. We inhibited the cell cycle in hBEC cultures following epithelial injury at the S phase using aphidicolin, an inhibitor of DNA synthesis, and at the G1 phase via selective CDK4/6 inhibition (PD0332991). While both compounds inhibited cycling as measured by EdU incorporation 11-14 days following epithelial injury (**Fig. 4F**), neither compound completely abrogated differentiation in regenerating cultures, as seen by immunostaining and qPCR (**Fig. 4G-H**) nor resulted in an upregulation of markers of the convergent programs associated with squamous epithelia (**Fig. 4H**). Aphidocolin-mediated S phase inhibition did result in a loss of multiciliated cells (**Fig. 4G-H**), potentially related to reported co-option of cell and centrosome cycles in multiciliation^46–49^. While further studies could help to determine this cell cycle phase-dependent mechanism, we may conclude that the loss of mucociliary differentiation induced by multiple ligands is not induced by reduced proliferation, and that proliferation is not a general requirement for normal luminal cell differentiation in ALI cultures regenerating after injury.

**Figure 4.**
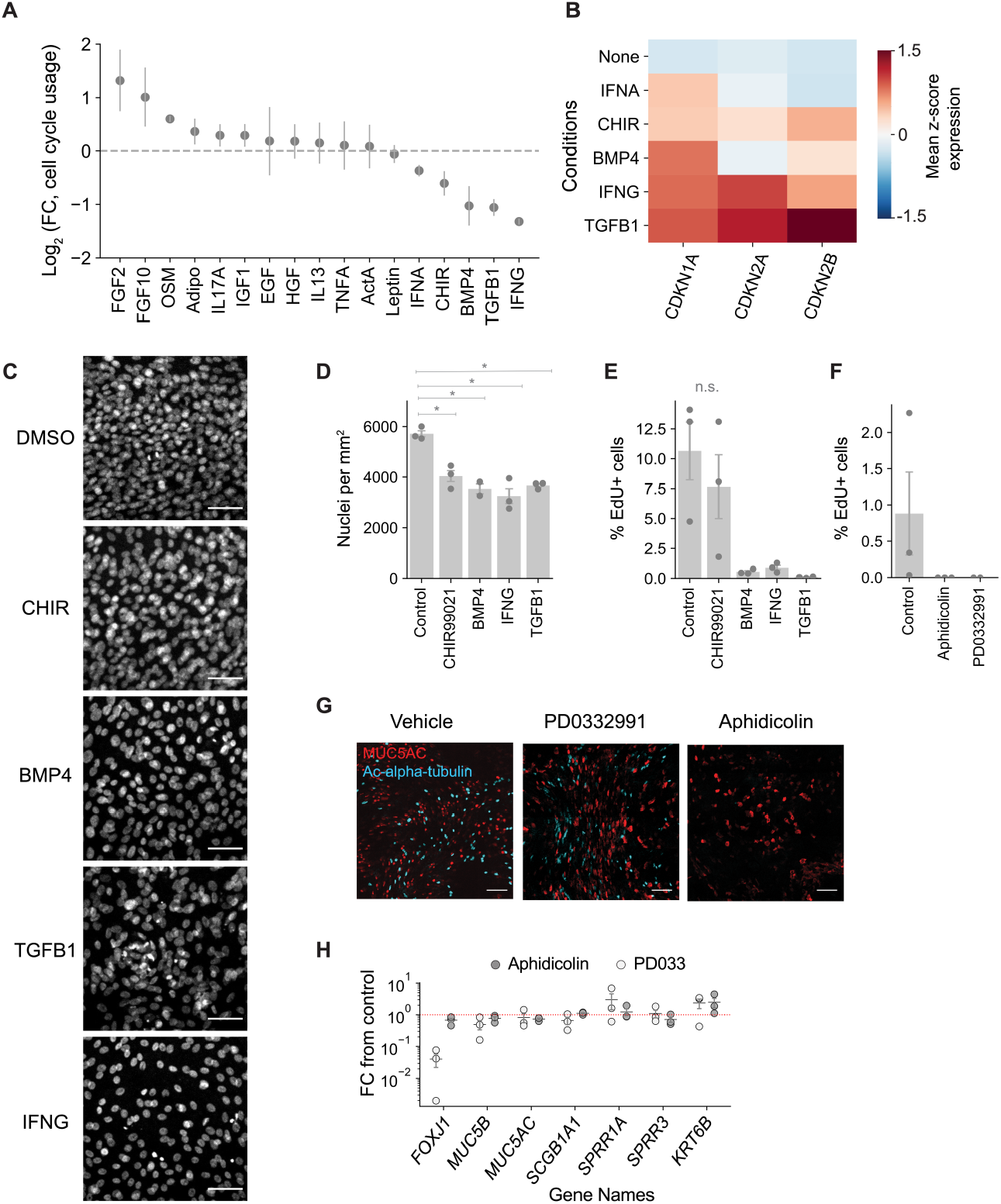
Cell cycle arrests during signaling-induced loss of airway epithelial mucociliary differentiation, but is not required for differentiation. **(A)** Fold change in cell cycle transcriptional program usage (defined in **Fig. S5A**) predicts a reduction in cell cycle in response to several signaling conditions. **(B)** Expression of cell cycle inhibitor genes in control, BMP4, CHIR, IFNG and TGFB1 induced cells. **(C)** Representative immunofluorescence images of whole-mount undifferentiated hBEC cultures treated with indicated cytokines and stained for DNA (Hoechst; white) indicate reduced cell densities in conditions showing reduced cell cycle programs and increased cell cycle inhibitor gene expression. Scale bars, 50 µm. **(D)** Quantification of areal cell density from panel C. Bars represent mean ± SEM, n = 3 HBEC donors (BMP4: n=2), calculated from 1 mm stitched images. *p≤0.05 by Wilcoxon rank-sum test. **(E,F)** Fraction of cells incorporating EdU after 48 hours of continuous EdU incubation **(E)** 48-hours following epithelial stripping and treatment with the indicated cytokines, or **(F)** 11-14 days following epithelial stripping and treatment with aphidicolin (2 ug/mL) and PD0332991 (100 nM). Bars represent mean ± SEM, n=3 HBEC donors. **(G)** Representative immuno-fluorescence images of whole-mount differentiated hBEC cultures treated with aphidicolin and PD0332991 and stained for MUC5AC (blue) and acetylated-alpha-tubulin (red). Scale bars, 100 µm. **(H)** Fold change of mRNA expression in differentiated HBEC cultures treated with aphidicolin/PD0332991 over untreated differentiated HBECs. Bars represent mean ± SEM, n=3 HBEC donors.

### Predicting signatures of signaling in human disease

Systematic maps of signaling in human tissues could help identify pathways that induce tissue disorganization in disease. This can be done by comparing the features of primary patient tissues to unique signatures associated with each pathway. Such features could be based on imaging, but transcription has many advantages in being scalable and offering multiple dimensions through which to identify signaling responses. Conventionally, transcriptional changes observed in disease have been studied by gene set enrichment analysis, but universal gene sets^50–52^ do not account for tissue-specific differences in signaling responses. We explored the use of transcriptional responses to signaling to generate hypotheses for signaling pathway activity in disease.

We developed a strategy to identify enrichment of the signaling-associated programs learnt from our data by gene signature enrichment^53^, here adapted to compare matched cell types from disease samples to their counterparts in control samples (**Fig. 5A**). We applied this strategy to published datasets of three lung diseases covering a total of 124 control or patient tissue samples from idiopathic pulmonary fibrosis (IPF), chronic obstructive pulmonary disease (COPD), and Covid-19 (**Fig. 5B**)^27–30^.

**Figure 5:**
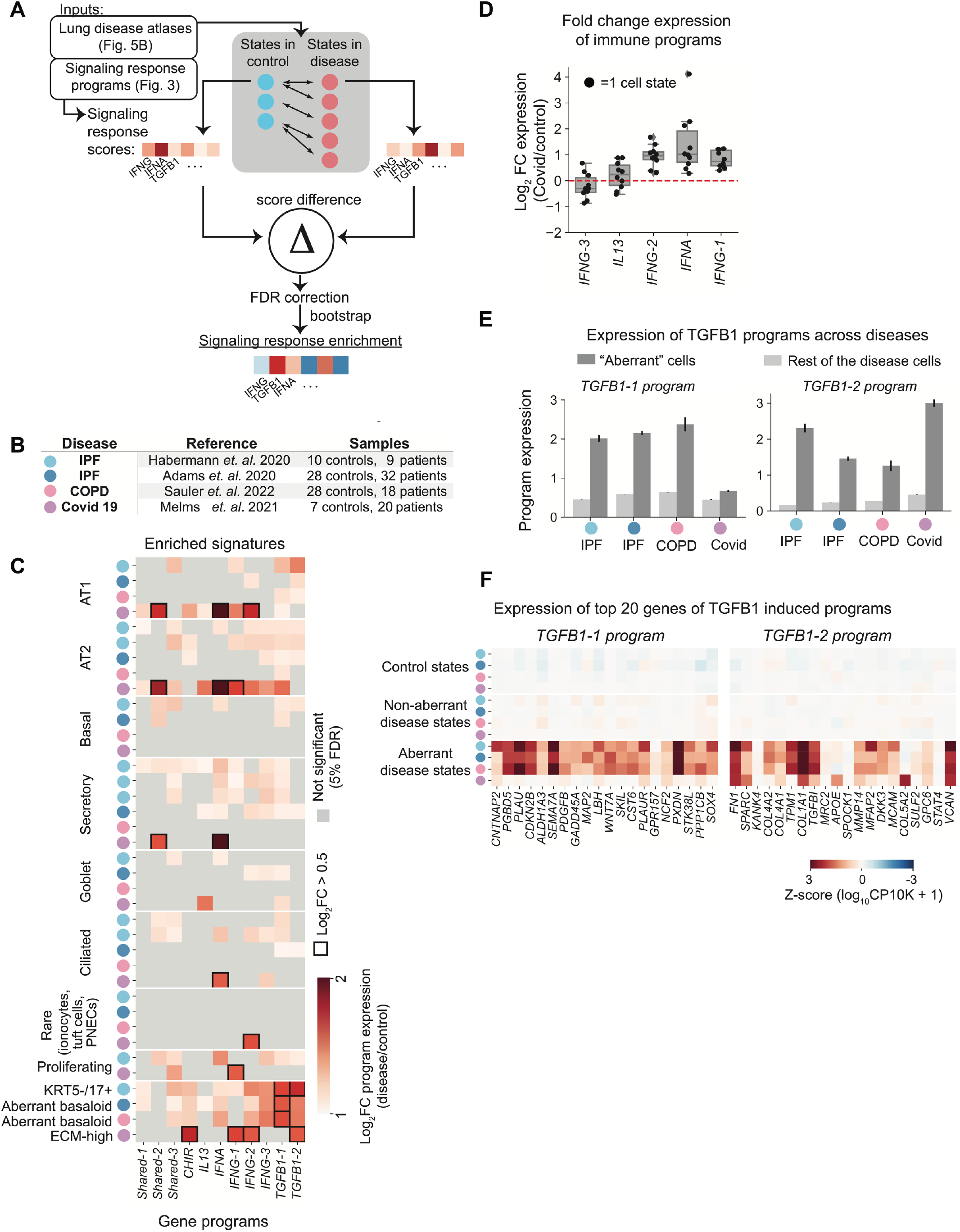
Evidence of direct signaling-induced states in lung disease atlases. **(A)** Schematic to infer signaling signatures in disease data by comparing matched cell states between disease and control samples. A signaling score is calculated using gene loadings from Fig. 3E for each cell state, and then compared by rank-sum testing with random downsampling of genes to ensure no single gene dominates the score. See Methods for detailed approach. **(B)** List of disease datasets used for the analyses. Each dataset is assigned a color that is used to represent these datasets in the subsequent figure panels. **(C)** Fold change increase in signaling program usage in disease versus control states across all diseases. Colors indicate dataset of origin for each cell state, as in B. **(D)** Fold change expression in immune signaling programs in Covid-19 cells. Each point represents a cell state. **(E)** Induction of TGFB1 signaling programs in cells identified as aberrant in the original papers, compared to other cells. Aberrant states include Krt17+/5-, aberrant basaloid, aberrant basaloid and ECM-high states in the 4 disease datasets respectively.

This strategy revealed a statistically significant enrichment of all eleven signaling-induced programs, relative to matched control samples, in a range of cell types across the four data sets (**Fig. 5C**). We highlight two observations that build confidence in the analysis. First, in the COVID-19 samples in particular, multiple cell types showed upregulation of inflammatory programs as expected during a viral response. However, in this disease not all immune programs were expressed: the *IFNA* and *IFNG-1/2* programs were upregulated across multiple cell types after COVID-19 infection, but *IL13* and *IFNG-3* programs were not (**Fig. 5D, S4**). These observations suggest that our map indeed captures specific epithelial responses to viral infection and predicts which inflammatory pathways are most active. Second, the IPF and COPD disease data sets studied here have reported the presence of an aberrant cell state high in KRT17 expression. These cells have been suggested to locally activate TGF-beta through their expression of integrin α_v_β_6_ subunits and through their localization directly lining myofibroblast foci^30,54,55^. In addition, analyses of these data suggested an enrichment of TGFB1 signaling responses across the entire data set based on Gene Ontology enrichment^50^, consistent with a central role for TGFB1 in fibrotic disease progression. We found here that the while TGFB1 programs were indeed broadly induced across airway cell types in IPF (**Fig. 5C**), the aberrant cells in these data sets expressed the two TGFB1-induced programs much more strongly than canonical cell types (**Fig. 5C,E**), and almost entirely recapitulated the program induced by TGFB1 in vitro (**Fig. 5F**). Notably, the gamut of genes enriched in this aberrant state including fibronectin (*FN1*), collagen 1A1 (*COL1A1*) and TGF-beta induced (*TGFBI*) arise uniquely from TGFB1 stimulation out of the pathways that we evaluated (**Fig. 3E, 5F**). This analysis also shows enrichment for other signaling pathway signatures in these aberrant cell state, with very similar patterns between the different IPF samples and COPD. By contrast, in COVID patient samples, an aberrant ‘ECM-high’ state has been reported. This state was enriched for only one of the two TGFB1 programs as compared to healthy lungs, and showed strong enrichment for an *IFNG* response that was absent in IPF and COPD. Thus, mapping signaling response in airway epithelium may provide a gateway to identifying direct cellular signaling responses in disease. The approaches used in this study can be expanded beyond airway epithelium to understand how signaling acts on other complex tissues, and drives the changes induced in disease.

## Discussion

In this study, we constructed a map of changes in cell type composition and cell type-specific gene expression in a regenerating culture of the human airway epithelium, in response to stimulation of 17 signaling pathways. We observed that the two principal axes of variation in cell type abundance after treatment recapitulate the primary forms of tissue metaplasia seen in diseases of the airway (**Fig. 2**): the loss of luminal differentiation, and goblet cell hyperplasia. However, the detailed changes in cell type frequencies and the gene expression programs induced by perturbation (**Fig. 3**) revealed far more granular phenotypes induced by signaling, including both convergent responses and unique signatures of several pathways evaluated here. In some cases, a single stimulus induced variable responses in different cells, as seen in the case of IFNG that induced three independent programs characterized by peak expression of either *IL1RL1*, or *CXCL11*, or *B2M*; and in the case of TGFB1 that induced an EMT response (*FN1*-hi) as well as a second program (*CDKN2B*-hi), and CHIR that induced both a convergent squamous-like program (*Shared-*1, high in *SPRR* genes) and a specific *KRT6C*-hi response (**Fig. 3, Table S6**). We showed that the near-total loss of control transcriptional programs in IFNG and TGFB1 is associated with changes in epithelial organization seen by loss of epithelial integrity. We further demonstrated that multiple perturbation programs associated with loss of normal differentiation induced cell cycle inhibition, but that the latter is not sufficient to arrest differentiation (**Fig. 4**).

These maps also offer an opportunity to investigate the regulation of rare cell types including the FOXI1+ pulmonary ionocyte, which have not so far been evaluated in almost any signaling context. Previously we showed a requirement for Notch signaling in ionocyte differentiation^5^, however, little is known about the role of other signaling pathways in maintenance and differentiation of this or other rare cell types. Here, we found that no signals clearly led to increased differentiation into ionocytes, PNECs or tuft cells, and indeed the proportion of these cells was maintained across multiple signaling conditions, suggesting that their differentiation may be under shared control not explored here. However multiple signals led to loss of this population alongside a global repression of luminal cell differentiation. The approaches utilized here could help to identify signaling conditions that modulate rare cell type abundance in the lung, with potential therapeutic relevance.

Together, these results offer a birds-eye view of the major axes by which the airway epithelium can be remodeled. They also provide a platform by which to interpret changes observed in scRNA-Seq atlases of human tissues. In the context of lung disease, our map established that a recently identified disease-specific (‘aberrant’) cell state directly recapitulates the expression response of airway basal cells to TGFB1. These cells have previously been suggested to locally activate TGFB1 through their expression of α_v_β_6_ integrin subunits. Our results support a view that these cells not only have a potential to activate TGFB ligands, but that they are directly induced by these ligands, a signature of a positive feedback loop in TGFB signaling^54,55^. This supports the hypothesis that lung fibrosis is sustained by a fibrotic cascade where abnormal signaling is perpetuated by the altered niche environment, and that epithelial cells may remodel their own niche^54,55^. We expect that this map will be useful to interpret further studies of lung disease, and to generate hypotheses for how signaling pathways maintain aberrant cells states in these diseases.

## Limitations of study

This study offers a strategy to define signaling responses during stem cell differentiation and identify these responses in disease. It has two types of limitations. First, it studies only the epithelium in isolation from its native environment, and lacking interactions with immune cells, fibroblasts, smooth muscle cells and extracellular matrix. For human tissues, the possibility to assay complete physiological responses is practically limited, but some of the responses identified here could be evaluated in animal models, albeit at lower throughput. Second, the number of pathways evaluated is not exhaustive, and we have not evaluated the response of the ligands over a range of concentrations or durations of exposure. We have also not evaluated their combinatorial effects, or the effects of inhibiting signaling pathways, or the contribution of secondary stimulation of one pathway by another by simultaneously activating and inhibiting pairs of pathways. As single cell analytical tools have advanced in the last few years, and allow for systematic sample multiplexing, one can now consider extending the map in these directions to define the range of signaling responses occurring in human tissue maintenance, regeneration, and disease.

## Acknowledgements

We are grateful for the support from the HMS Single Cell Core, and to Yuheng Lu for support with sample genotype demultiplexing. We thank Lindsey Plasschaert, Cathy Quigley, David Rowlands at Novartis Institutes for BioMedical Research (NIBR) and members of the Klein lab for their comments. This work was supported by a Cystic Fibrosis Foundation postdoctoral fellowship to Katherine McCauley, NIH grants R33CA212697 and R01HD096755 and by an SRA from NIBR to AMK.

## Materials and Methods

For detailed methods, please refer to the accompanying Supplementary Information.

