## Supplementary material for "A map of signaling responses in the human airway epithelium": Materials and Methods

### Contact for Reagent and Resource Sharing

### Experimental Models and Subject Details

#### Selection of receptors and pathways

*Data sources and pre-processing.* A list of genes encoding receptors with at least one known ligand supported by published data were obtained from an annotated ligand-receptor database (Table S3 of Ramilowski et al.[31]). Genes were considered if their receptor was paired with ligands and annotated as "known" and "literature-supported" in column "Pair.Evidence" of the database. The resulting list of receptor gene symbols is denoted . Selection of receptors from this list was performed on scRNA-Seq data obtained from Plasschaert et al.[5], GEO accession GSE102580. Cell-by-gene counts matrices were obtained from files *GSE102580_filtered_normalized_counts_human.tsv* and *GSE102580_filtered_normalized_counts_human_viral_transduction.tsv*, and cell type annotations were obtained from files *GSE102580_meta_GSE102580_filtered_counts_human.tsv* and *GSE102580_meta_filtered_counts_human_viral_transduction.tsv*. Cells with annotation "Basal" were pooled from both data sets, averaged across all cells, and then normalized to units of transcripts per million (TPM) to give the mean expression of gene *k* in basal cells, . For comparison to other tissues, data were downloaded from the GTex Portal Bulk RNASeq database (gtexportal.org/home/datasets; file *GTEx_Analysis_2017-06-05_v8_RNASeQCv1.1.9_gene_median_tpm.gct.gz*). Data was normalized to TPM, and the average and standard deviation expression for each gene *k* across all tissues was calculated. A standardized expression of each gene *k* in basal cells relative to GTEx tissues was then calculated as /.

*Receptor-ligand selection and prioritization.* Genes were then filtered according to three criteria: (1) the gene is in the list ; (2) it has minimal expression ; and (3) it is expressed at average or higher levels relative to other tissues . Receptors that passed these three filtering steps were then categorized based on known literature function into categories "Developmental signaling," "Immune signaling," "Hormone receptors," "Matrix interactions," and "Other," with the latter including gene products implicated in cell-cell adhesion, cell motility, and proliferation as well as decoy receptors and death receptors. "Developmental signaling," "Immune signaling," and "Hormone receptors" were selected. Ligands cognate to the selected 31 receptors were selected from Table S3 of Ramilowski et al.[31] and published literature. Where multiple ligands were available, those binding to multiple receptors were prioritized.

#### hBEC Culture and Cytokine Treatments

Primary human bronchial epithelial cells (HBECs) from normal donors were obtained from Lonza (Cat. # CC-2540) and were expanded for two passages with Growth Medium (BEGM supplemented with one SingleQuots kit, Lonza Cat. # CC-3171) in uncoated T75 flasks (Corning Cat. # 430641U). After expansion, cells were seeded on uncoated 12-well TransWell (Corning Cat. # 3460) plates at a density of 1e6 cells per plate and expanded for 7 days in Differentiation Medium (50:50 BEGM (Lonza Cat. # CC-3171)/high-glucose DMEM (ThermoFisher Cat. # 11965175) supplemented with one SingleQuots kit with no T3 and final retinoic acid concentration of 50 nM, Lonza # CC-4175). After 7 days, media was removed from the apical surface and cells were differentiated at air-liquid interface for an additional 30-45 days with media replaced every 48-72 hours. Cells were then washed 3X with warm 1XS PBS and treated with MEM (ThermoFisher Cat. # 10370088, 500 µL apical, 1 mL basal) for 1 hour at 37°C to remove differentiated luminal cells. Following incubation, luminal cells were removed by pipetting gently up and down 3-5X before removing apical media and washing with 500 µL MEM in apical chamber[40]. Cytokines were diluted in Differentiation Medium at concentrations typically 10-100X higher than reported IC50 doses, as indicated in Supplemental Figure 1B. Cells were cultured for an additional 14 days, replacing media every 48-72 hours.

#### Single-cell dissociation and capture for single cell sequencing

HBECs were collected 14 days post-injury by treating with 0.05% Trypsin-EDTA (prepared by dilution of 0.25% Trypsin-EDTA (ThermoFisher Cat. # 25200056) with calcium/magnesium-free PBS), for 20 minutes at 37 °C. Trypsin dissociation was halted with Trypsin Neutralizer Solution (ThermoFisher Cat. # R002100) at 1:1 dilution and cells were filtered through a 40 µm strainer and pelleted by centrifugation at 300x g for 5 minutes. Cells were resuspended in phosphate buffered saline (ThermoFisher Cat. # 10010072) with 0.1% bovine serum albumin (ThermoFisher Cat. # B14) and counted on a Countess Cell Counter (ThermoFisher) to determine cell number and viability. For cell counting, 10 µL of cell suspension was mixed with 10 µL of Trypan Blue Solution (ThermoFisher Cat. # 15250061) and 10 µL of that 1:1 mix was loaded into Countess chip and read with default program. Viable cell numbers were used to determine cell concentration. Optiprep Density Gradient Medium (Sigma Cat. # D1556) was added to achieve a final concentration of 15% Optiprep and 120-150,000 cells/ml. Cells were run in two separate experiments: donor #429581 was collected individually; donors #221175 and #323353 were pooled and captured simultaneously.

scRNA-Seq was performed using inDrops[56], following previously described protocols[57]. Briefly, single cells and hydrogel beads tagged with barcoding primers for reverse transcription (RT) were encapsulated together in 3-4 nL droplets. RT was carried out in droplets in an emulsion. The emulsion was broken for library production consisting of (i) second strand synthesis, (ii) in vitro transcription for linear amplification, (iii) RNA fragmentation, (iv) RT, and (v) amplification by polymerase chain reaction (PCR). The resulting libraries were sequenced on NovaSeq and NextSeq Illumina platforms in paired-end mode at length of 2x76 base pairs and converted into FASTQ files using standard Illumina pipelines. Data was processed into BAM files using published pipelines[57].

### Single-cell RNA sequencing read processing, data filtering and normalization

Gene expression counts matrix was generated from raw FASTQ files using indrop.py pipeline[56] (https://github.com/indrops) using human genome assembly GRCh38/86 (genome assembly/ENSEMBL release). For donor lots #221175 and #323353 where cells were pooled prior to droplet capture (see above), the cell barcodes were then assigned to each donor respectively by their SNP profiles using the scSplit package[58]. To retain high quality transcriptomes, total count and a mitochondrial count filters were applied. Transcriptomes with more than *N* total UMI counts and less than 40% of counts coming from mitochondrial genes were retained. The UMI threshold *N* was chosen separately for each library within the range 500-800 after manual inspection of the UMI/cell distribution per library. The final threshold values used can be determined from the minimal total UMI counts/cell per library in metadata file `meta_data_unfiltered.tsv` on GEO. For mitochondrial gene count calculations, genes with a gene symbol starting with "*MT-*" defined the list of mitochondrial genes. Subsequently, scRNA-Seq data were processed in Python version 3.7+ using the scanpy (sc) package (version 1.6+)[59]. For normalization, genes expressed in less than 3 cells and with less than 6 total counts across all cells were filtered out using the function and parameters - `sc.pp.filter_genes(min_cells=3, min_counts=6)` and then cell counts were normalized to total 10,000 counts for each cell using `sc.pp.normalize_per_cell(counts_per_cell_after=1e4)` to define as the expression of gene *j* in cell *i* in units of counts per 10,000 (CP10K). Counts were log10-normalized using the function `sc.pp.log` and default parameters).

### Dimensionality reduction and with donor integration of single cell data

Dimensionality reduction was carried out twice – one for untreated cells only (**Fig. 1C,E**), and one for the complete data set (**Fig. 1D** and all subsequent analyses). In each of the two cases, to reduce dimensionality, the top 4000 highly variable genes were identified using `sc.pp.highly_variable_genes(flavor='cell_ranger', n_top_genes=4000)`, and counts were z-scaled using `sc.pp.scale` to give where is the set of cells being analyzed (untreated cells or all cells), and are respectively the mean and variance over cells . For untreated cells, this step was followed by Principal Component (PC) analysis with number of PCs = 50, and a k-nearest graph was constructed with correction for donor variation via the bbknn algorithm (batch balanced k-nearest neighbors)[33] (version = 1.31, function: `bbknn.bbknn(batch = 'assigned_donor', neighbors_within_batch = 10 )`). For the full data set, we tested several methods for donor batch correction, and found upon visual inspection that Harmony (version 0.05) adequately preserved within-donor variation. We constructed a batch corrected embedding with Harmony using `harmony.run_harmony(vars_use='experiment')` function with PC coordinates as input. We then constructed a k-nearest neighbor graph with k=20 nearest neighbors in the donor-corrected embedding. In both cases, the embedded data was visualized into two dimensions using a UMAP projection using function: `sc.tl.umap(random_state = 0)` (**Figs. 1C,D**).

### Annotation of untreated hBEC sample scRNA-Seq data

After performing dimensionality reduction and k-nearest neighbor graph construction (above), the untreated data was clustered using Leiden clustering[60] (function: `sc.tl.leiden`, default parameters). The clusters were manually annotated as: basal, KRT13 high, KRT17 high, secretory, multiciliated or a cluster of "rare cells" including PNECs, tuft cells and ionocytes. Annotation was determined according to the mean expression of cell type-specific marker genes shown in **Fig. 1F**. The "rare cells" cluster was then subclustered and cells in the resulting clusters were annotated as ionocytes, tuft and PNEC cells after inspection for marker genes shown in **Fig. 1F**.

### Composite gene scores

The subsequent sections make use of composite gene scores as defined here. For a set of cells , we define the z-standardized gene expression for gene *j* in cell *i* as (see above for z-standardization). We then define the knn-smoothed value of to be , where is the set of *k* nearest-neighbors of cell *i* in the 50-dimensional PC space (see above). For a gene set , we define a composite gene score for cell *i* as the average of the smoothed z-standardized expression of the genes in the set, = .

### Cell type classification in treated samples

To classify perturbed cells, a logistic regression classifier was trained on annotations of untreated data. Python's machine learning package `sklearn` (version: 0.24.2+) was used with function and parameters: `sklearn.linear_model.LogisticRegression(c_param=1, n_iterations = 1000)`.The 50 PC values learnt from untreated cells (see above) were used as features for classification. The annotations "basal", "KRT13-high" and "KRT17-high" were merged into one class ("basal"). The gene expression values for treated cells was standardized to the control samples and then projected onto the PCs of untreated cells. The trained classifier applied to label each treated cell. Two subsequent steps were then carried out to (1) relabel the cells classified as "secretory" as either

“goblet” or “club”, and (2) identify cells with low-confidence assignment and relabel these as “indeterminate”. The two steps are as follows:
*Classifying goblet vs club cells.* A composite “goblet gene score” =10) was calculated with . Cells classified as secretory were sub-labeled as “goblet” if , and “club” otherwise.

*Declassifying indeterminate cells*. For multiciliated cells, ionocytes, tuft cells and PNECs, a composite gene score =10) was calculated for each cell using the following gene sets:
Multiciliated: $S_{multiciliated}$ = *FOXJ1, C9orf24, RSPH1, TPPP3* and *SNTN*
Ionocyte: $S_{ionocyte}$ = *FOXI1, TMPRSS11E, STAP1, CFTR* and *PDE1C*
Tuft: $S_{tuft}$ = *POU2F3, LRMP, NREP, CRYM* and *HOTAIRM1*
PNEC: $S_{PNEC}$ = *ASCL1, SST, GRP, CALCA* and *CHGA*
Let be the annotated cell type label assigned to cell *i* using the logistic regression classifier. A cell was re-labeled as “indeterminate” if it expressed its label-associated marker gene score below a threshold value . The threshold value was manually determined for each annotation by comparing the distribution of between untreated cells with annotation to all cells with other annotations . The two distributions for each cell type are shown in **Fig S2A**. Cells labelled as “indeterminate” were excluded from downstream cell type specific analyses in **Figs. 2-3**.
**Fig. S2A** also shows comparable gene scores for basal and secretory cells, using the gene scores:
Basal: $S_{basal}$ = *KRT5, KRT13, S100A2, KRT14* and *CSTA*
Secretory: $S_{sec}$ = *MUC5B, SCGB3A1, VMO1, BPIFA1* and *BPIFB1*

### Fraction control neighbor analyses (Fig. 1G)

For each donor separately, let be the set of untreated cell transcriptomes in the data set post-filter, and let be the set of cells from treatment *t.* For each cell *i*, the set of 100 nearest neighbor transcriptomes was determined by Euclidean distance on the Harmony-corrected principal component space (see section “Dimensionality reduction and with donor integration of single cell data”). Then, is the total number of control cells among the 100-nearest-cell neighborhood of cells in treatment *t*, and is the respective number of cells from treatment *t*. Defining the control fraction , a null expectation for is ) where are the total number of cell transcriptomes in each set respectively. This null represents the assumption that treatment does not alter cell states relative to untreated controls. The observed/expected ratio is . This quantity was calculated for each donor and treatment condition. The mean and SEM across replicate donors are plotted in **Fig. 1G.**

### Gene expression heatmaps

For all gene expression heatmaps in the paper, gene expression was quantified as [z-score of $\log_{10}$(CP10k + 1)], or as z-score of CP10k as specified in figure captions. For reporting gene expression of single cells rather than cluster means (**Fig. 2B**, **2K**), we plot graph-smoothed gene expression as defined above. For **Fig. 2B**, the cells have been ordered first according to their cell type annotations and then within cell type, they have been ordered in decreasing order of mean expression of cell type specific marker genes. For **Fig. 2K**, goblet cells are first separated by condition and then ordered by decreasing order of *MUC5AC* expression.

### Testing and visualizing changes in cell type abundance (Fig. S2B, Table S3)

For p-values in **Table S3,** Fisher’s exact test was used to evaluate statistical significance in the changes in cell type frequency between treated and control samples separately for each donor. The resulting p-values across donors were combined by Fisher’s chi-squared method and controlled for false discovery rate by the Benjamini-Hochberg method. For **Fig**. **S2B**, we plot for each donor, where corresponds to the frequency of cell type in the *t*-th treatment condition, and is the respective cell type frequency for the untreated control. Where , we note “n.d.” (not detected). Line averages in **Fig. S2B** and bar chart values in **Figs. 2F,H** plot where is an average over donors for which each treatment was carried out. For principal component (PC) analysis of changes in cell type abundance (**Figs. 2C,D**), we calculated , where e=0.1% is a pseudocount. PC analysis was then carried out on the scaled values using Python `sklearn.decomposition.PCA` (sklearn package version 0.24.2+), where and are the mean and variance of each cell type *l* over all treatments *t*.

### Differential gene expression per cell type

Tests for differential gene expression were performed between cells from each treatment and untreated sample, for cells with matched cell type labels – basal, secretory (club + goblet) and rare cells (ionocytes + tuft + PNEC cells). Normalized filtered gene expression counts (see above) were used for testing. Statistical significance was calculated by the Wilcoxon rank sum test python `scipy` (version 1.9.3), with multiple hypothesis correction controlling for false discovery rate using Benjamini and Hochberg’s method (python `statsmodels` package, version 0.13+). Gene fold-changes in expression per cell type, reported in **Table S4**, were calculated with a pseudo-count correction where is the set of cells with label and treatment . The number of genes with >1 and FDR < 0.05 are reported in **Fig. 3A**.

### Treatment program factorization using cNMF

Gene expression programs were learnt by consensus non-negative matrix factorization (cNMF)[38,61] using the Python package (`cNMF`, version 1.1) with parameters: number of components k = 20, percentage of replicates used as nearest neighbors for outlier detection = 30%, and local density threshold for defining outliers = 0.2. Usage and program matrices are provided in **Tables S5, S6**. The resulting program usages, , are normalized such that for programs *j* in each cell . The gene weights are normalized such that for program *j* over all genes (shown in **Fig. 3E**, **Table S6**).

### Visualization of cNMF program usages (Figs. 3C,D,F S3A,C)

For visualization of treatment-specific usages in **Figs. 3C, S3A**, the program usages were averaged across all cells and then all donors for each condition: for every donor *i,* program *j* and treatment condition *t*, program usage where and are respectively the set of cells from treatment *t* and donor *i*, and . The average usages were then averaged across donors, , and scaled for plotting in **Figs. 3C, S3A** (top). For **Fig. S3A** (bottom), the fold-changes in usage shown for each treatment and each program are , were corresponds to the untreated controls. For visualization of cell type-specific usages of control programs (**Figs. 3D, S3C**), mean usages were similarly calculated, , but now using the following cell sets . For **S3C**, we plot only for cells from the untreated control samples, . For shared perturbation programs (**Fig. 3D** first panel), we plot only for cells from the treated samples, . And for perturbation programs unique to specific signals (**Fig. 3D,** except first panel), we plot mean usage of only the cells from the respective signaling perturbation, .

### Aggregate control program usage

To plot usage of control in specific conditions for different cell types (**Fig. 3F**), usages of all control programs was summed for each cell, . The median value of across all cells of the indicated cell types (basal, secretory, multiciliated and rare cells) in the indicated treatment conditions was plotted. BMP4 and IFNG have ≤1 rare cells and hence are not included in the rare cell plot (**Fig. 3F**, last panel). Aggregate perturbed program usage in each cell is equal to 1-.

### Enrichment of signaling programs in disease (Fig. 5)

Human scRNA-Seq data sets consist of an expression matrix (units of CP10K) for cell and gene , with annotations representing a set of state labels for each cell. To define signaling responses, we used gene sets each consisting of the *M* genes with the largest cNMF weights for each program *j* (defined above). To calculate a score from , we define a sparsified weight matrix , with normalization factor chosen such that . We further define a (randomly) downsampled sparsified weight matrix with being a random sample of *m* integers between 1 and M without repetition. Hereafter we use *M*=20 and m=7, and for brevity omit arguments for . The signature score for program *j* in a single cell is then calculated as a weighted sum , and the randomly-downsampled signature score is . To avoid any single gene dominating the signature scores, we bootstrap a distribution of by 100 instantiations of , and obtain the median score ].

We calculated a bootstrapped signature score for every cell in each of the data sets shown in **Fig. 5B,** for each cNMF program shown in **Fig. 5C**. A one-tailed Wilcoxon rank-sum test was then carried out between the values for cells annotated with the same label between disease and healthy donor samples to find programs that are significantly upregulated in diseased cells. As an exception, for the IPF and COPD "aberrant basaloid" state and "*KRT17*+/*KRT*5-" state there are less than 0.15% corresponding healthy cells; for these states we instead carried out comparison to cells annotated as "basal" in healthy donor samples. The resulting p-values were then controlled by the Benjamini-Hochberg procedure and significance was determined at 5% FDR. The magnitude of signaling response was calculated for each labeled state as where represent the set of cells with matched state label in the disesase (*d*) and healthy (*h*) donor samples respectively (**Fig. 5C**).

### Immunostaining of ALI cultures

For immunofluorescence on cross-sections of TransWell cultures (**Fig. 2**), membranes were fixed in 10% neutral-buffered formalin overnight at room temperature (12-24 hours), then transferred to 70% ethanol solution, excised by scalpel and embedded in paraffin blocks using standard dehydration and processing approaches. 5 μm sections were cut on a tissue microtome, mounted on 1.0 mm glass slides, dried and de-paraffinized following standard protocols.

Whole mount cells were fixed in fresh 4% paraformaldehyde (PFA, diluted from 16% stock in 1XPBS) for 1 hour at room temperature, washed 3 x 20 minutes with PBS at room temperature, and stored at 4°C for no more than 14 days until staining.

Both sections and whole-mount cultures were incubated for 1 hour at room temperature in immunofluorescence buffer (IF Buffer, 130 mM NaCl, 7 mM $Na_2HPO_4$, 3.5 mM $NaH_2PO_4$, 7.7 mM $NaN_3$, 0.1% bovine serum albumin, 0.2% Triton X-100, and 0.05% Tween- 20) supplemented with 10% normal goat serum. Primary antibody was incubated overnight at 4°C, washed for 3 x 20 minutes with IF buffer, and secondary antibody plus 1:5000 Hoechst 33342 was incubated for 1 hour at room temperature. Cells were washed 3 x 20 minutes at room temperature in 1X PBS then mounted with coverslips with ProLong Diamond Antifade Reagent (ThermoFisher). F-Actin staining was performed by incubating with phalloidin dye (ThermoFisher) diluted 1:300 in 1XPBS for 30 minutes at room temperature. The stained samples were imaged on a confocal microscope: Axiovert 200 microscope (Carl Zeiss) with Yokogawa CSU-X1 spinning disc head and an Evolve 512 electron-multiplying charge-coupled device camera (Photometrics). Raw images were processed by adjusting brightness and contrast, merging channels, adding scale bars, and adding false-color, using ImageJ. Antibodies used for immunofluorescence are detailed in the Reagents and Resources Table.

### Permeability assay

Differentiated HBEC cultures were prepared from n=1-3 donors as above (Lonza, donors 134626, 646466, 627466). Following 1 week of differentiation, cultures were stripped as described previously and differentiated for an additional two weeks in media supplemented with CHIR99021, rhTGFB1, rhBMP4, IFNG, and IFNA, at concentrations listed **Fig. S1B**.

Cells were washed 3 x with HBSS and incubated for 1 hour at 37°C in HBSS (basal chamber) and HBSS + 0.1 mg/mL Lucifer Yellow (apical chamber, Sigma). 150 μL of media was transferred from the basal chamber to a 96-well assay plate (Greiner) and measured in a spectrofluorometer (Clariostar, BMG LabTech; excitation=485 nm, emission=535 nm). Data were normalized to a blank well containing HBSS and the resulting value from the control well was set to 1.

#### EdU measurement in cytokine-treated cultures

Differentiated HBEC cultures were prepared from n=3 donors as above (Lonza donor #s 134626, 646466, 627466). Following 1 week of differentiation, cultures were stripped as described previously and cultured for 48 hours in media supplemented with CHIR99021, rhTGFB1, rhBMP4, or IFNG, at concentrations listed in Supplemental Figure 1B. After 48 hours, cells were treated with 2 μM ClickIt EdU labeling reagents (ThermoFisher) diluted in culture media for an additional 48 hours at 37°C. EdU incorporation was measured by flow cytometry on a Cytoflex cytometer (Beckman).

#### Nuclear quantification in cytokine-treated cultures

Differentiated HBEC cultures were prepared from n=3 donors as above (Lonza donors 134626, 646466, 627466). Following 1 week of differentiation, cultures were stripped as described previously and cultured for 14 days in media supplemented with CHIR99021, rhTGFB1, rhBMP4, or IFNG, at concentrations listed in **Fig. S1B**. Cells were fixed with 4% PFA and nuclei were stained with Hoechst dye (1:5000 dilution, Thermo Fisher) at room temperature for 1 hour. Membranes were excised using 0.8 cm biopsy punch and mounted on 1.0 mm glass slides with glass coverslips (#1, 30x22 mm at 13 mm thickness) in ProLong Diamond Mounting reagent. One BMP4-treated sample was lost during mounting, so BMP4 data is presented as n=2. For each condition, nuclei were imaged on a confocal microscope using automated built-in stitching software with starting point at center of mounted membrane and 10% overlap between images (Zen Blue Microscopy software; Carl Zeiss) to generate 3260 x 3260 μM 20x images. Raw images were processed and analyzed using ImageJ.
(i) Loaded in full stitched image and ran auto-adjust brightness/contrast (built-in ImageJ function)
(ii) Automatically applied built-in ImageJ functions for smoothing, thresholding (Huang fuzzy thresholding method [62]), and watershedding. This approach is used to automatically distinguish closely-grouped nuclei to allow counting of individual cells.
(iii) Cropped image to 1 $mm^2$ to eliminate edge effect variability across samples
(iv) Calculated centroid measurements of minimum size 1 $pixel^2$ (1 centroid = 1 cell)

#### Effect of cell cycle inhibitors on differentiation

Differentiated HBEC cultures were prepared from n=3 donors as above. Cells were pre-treated for 48 hours with 2 μg/mL aphidicolin (Sigma) or 1 μM PD0332991 (Sigma), then stripped as described above. 11-14 days post-stripping, cells were treated with 2 μM ClickIt EdU labeling reagents (ThermoFisher) for 48 hours or with 10 μg/mL EdU for 4 hours, and EdU incorporation was measured by flow cytometry on a Cytoflex cytometer (Beckman). Cells were harvested for immunofluorescence by fixation with 4% PFA and stained as above with primary antibodies against MUC5AC and acetylated alpha-tubulin. Antibody information can be found in Reagents and Resources Table.

#### Reverse Transcriptase Quantitative RT Polymerase Chain Reaction (qRT-PCR)

Cells were lysed with RLT Buffer (RNEasy kit, Qiagen), RNA was extracted following RNA was extracted following manufacturer's protocol, and 1 ug of RNA was transcribed to cDNA using reverse transcription reagents (High-Capacity RNA-to-cDNA kit, Thermo). cDNA was diluted between 1:4 and 1:6 and 4 μL of cDNA was added to each 10 μL TaqMan Fast Universal PCR Master Mix (ThermoFisher) qPCR reaction. Gene expression was detected using TaqMan (ThermoFisher) probes (gene-specific primer information in Methods Table). Relative gene expression for each sample was calculated using the $2^{(-\Delta\Delta CT)}$ method by normalizing the cycle number (Ct) for each sample to an 18S control and to a control sample, where baseline of control was defined as fold change = 1. TaqMan assay probes used are detailed in Reagents and Resources Table.

### Reagents and Resources Table

| REAGENT or RESOURCE | SOURCE | IDENTIFIER |
|---|---|---|
| **Antibodies** | | |
| Mouse monoclonal to acetylated alpha-tubulin (clone 6-11B-1) | Millipore Sigma | T6793 |

| | | |
|---|---|---|
| Mouse monoclonal to MUC5AC (clone 45M1) | ThermoFisher | MS-145-P0 |
| Purified anti-keratin 5 polyclonal chicken antibody | BioLegend | 905903 |
| Mouse monoclonal to MUC5B (clone 5B19-2E) | ThermoFisher | 37-7400 |
| Goat anti-Mouse IgG2b Cross-Adsorbed Secondary Antibody, Alexa Fluor 647 | ThermoFisher | A-21242 |
| Goat anti-Mouse IgG1 Cross-Adsorbed Secondary Antibody, Alexa Fluor 546 | ThermoFisher | A-21123 |
| Goat anti-Mouse IgG2b Cross-Adsorbed Secondary Antibody, Alexa Fluor 546 | ThermoFisher | A-21144 |
| Goat anti-Mouse IgG1 Cross-Adsorbed Secondary Antibody, Alexa Fluor 488 | ThermoFisher | A-21121 |
| Goat anti-Chicken IgY, | ThermoFisher | A-21449 |
| Alexa Fluor 647 Phalloidin | ThermoFisher | A22287 |
| **Biological samples** | | |
| Human bronchial epithelial cells (donor numbers 429581, 221175, 323353, 134626, 646466, 627466) | Lonza | CC-2540 |
| Chemicals, peptides, and recombinant proteins | | |
| Hoechst | ThermoFisher | H3570 |
| Recombinant human Activin A | R&D Systems | 338-AC-010 |
| Recombinant human BMP4 | Peprotech | 120-05 |
| Recombinant human EGF | ThermoFisher | PHG0311 |
| Recombinant human FGF10 | Sino Biological | 10573-HNAE |
| Recombinant human FGF2 | ThermoFisher | PHC9534 |
| CHIR99021 | Stem Cell Technologies | 72052 |
| Recombinant human HGF | ThermoFisher | PHG0254 |
| Human IFN-Alpha (alpha 2A) | PBL Assay Science | 11100 |
| Recombinant human IFN-Gamma | Peprotech | 300-02 |
| Recombinant human IL-13 | Peprotech | 200-13 |
| Recombinant human IL-17A | ThermoFisher | PHC9714 |
| Recombinant human Oncostatin M | Peprotech | 300-10 |
| Recombinant human TNF-alpha | Peprotech | 300-01A |
| Recombinant human Adiponectin | Peprotech | 450-24 |
| Recombinant human Leptin | R&D Systems | 398-LP |
| Trypan Blue Solution | ThermoFisher | 15250061 |
| Triton X-100 | Millipore Sigma | X-100 |
| Tween-20 | Millipore Sigma | P2287 |
| 16% Paraformaldehyde | Fisher Scientific | 50-980-487 |
| Lucifer Yellow CH dipotassium salt | Millipore Sigma | L0144 |
| Aphidicolin from Nigrospora sphaerica | Millipore Sigma | A0781 |
| Palbociclib (PD0332991) | Millipore Sigma | PZ0383 |
| ProLong Diamond AntiFade Mountant | ThermoFisher | P36965 |
| OptiPrep Density Gradient Medium | Millipore Sigma | D1556 |
| Formalin solution, neutral buffered | Millipore Sigma | HT501128 |
| Normal goat serum | ThermoFisher | 38172 |
| **Critical commercial assays** | | |
| Click-iT EdU Pacific Blue Flow Cytometry Assay Kit | ThermoFisher | C10418 |
| High-Capacity RNA-to-cDNA Kit | ThermoFisher | 4388950 |
| Taqman assay: *FOXJ1* | ThermoFisher | Cat. # 4331182, assay ID Hs00230964_m1 |
| Taqman assay: *MUC5B* | ThermoFisher | Cat. # 4331182, assay ID Hs00861595_m1 |
| Taqman assay: *MUC5AC* | ThermoFisher | Cat. # 4331182, assay ID Hs01365616_m1 |
| Taqman assay: *SCGB1A1* | ThermoFisher | Cat. # 4331182, assay ID Hs00171092_m1 |
| Taqman assay: *SPRR1A* | ThermoFisher | Cat. # 4331182, assay ID Hs00954595_s1 |
| Taqman assay: *SPRR3* | ThermoFisher | Cat. # 4331182, assay ID Hs01878180_s1 |

| | | |
|---|---|---|
| Taqman assay: *KRT6B* | ThermoFisher | Cat. # 4331182, assay ID Hs00749101_s1 |
| TaqMan assay: *18S ribosomal RNA* | ThermoFisher | Cat. # 4448481, assay ID Hs03928985_g1 |
| TaqMan Fast Universal PCR Master Mix (2X), no AmpErase UNG | ThermoFisher | 4366072 |
| RNEasy Plus Mini Kit | Qiagen | 74134 |
| **Software and algorithms** | | |
| ImageJ | National Institutes of Health | https://Imagej.nih.gov/ij/ |
| Indrop.py pipeline | Zilionis et al. (2017) | https://github.com/swolock/indrops |
| Zen Blue | Carl Zeiss | https://www.zeiss.com/microscopy/en/products/software/zeiss-zen.html |
| scSplit | Xu et al. (2019)[58] | https://github.com/jon-xu/scSplit |
| Scanpy | Wolf et al. (2018)[59] | https://scanpy.readthedocs.io/en/stable/index.html |
| cNMF | Kotliar et al. (2019)[38] | https://github.com/dylkot/cNMF |
| Python 3.7 or above | Anaconda | https://www.python.org/ |
| **Other** | | |
| BEGM Bronchial Epithelial Cell Growth Medium BulletKit | Lonza | CC-3171 |
| Bronchial Epithelial SingleQuots Kit | Lonza | CC-4175 |
| 96-well F-bottom black assay plate | Greiner | 655076 |
| 12 mm Transwell with 0.4 µm Pore Polyester Membrane Insert, Sterile | Corning | 3460 |
| 75cm² U-Shaped Canted Neck Cell Culture Flask with Vent Cap | Corning | 430641U |
| Micro Cover Glasses, Rectangular, no. 1 | VWR International | 48393-026 |
| Frosted Micro Slides | VWR International | 48312-004 |
| Trypsin-EDTA (0.25%) | ThermoFisher | 25200056 |
| Phosphate buffered saline, pH 7.4 | ThermoFisher | 10010072 |
| Fetal bovine serum | ThermoFisher | 16000044 |
| Hank's Balanced Salt Solution | ThermoFisher | 14025092 |
| Bovine serum albumin | ThermoFisher | B14 |
| Trypsin Neutralizer Solution | ThermoFisher | R002100 |
| MEM | ThermoFisher | 10370088 |
| DMEM, high glucose | ThermoFisher | 11965175 |
