## Supplemental Figures and Tables 1-2 for "A map of signaling responses in the human airway epithelium"

### Supplementary Figures

A

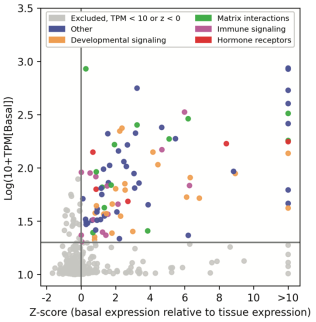


B

| Category | Ligand/ Molecule | Dose |
|---|---|---|
| Developmental | Activin A | 10 ng/mL |
| Developmental | BMP4 | 10 ng/mL |
| Developmental | EGF | 20 ng/mL |
| Developmental | FGF10 | 100 ng/mL |
| Developmental | FGF2 | 10 ng/mL |
| Developmental | CHIR99021 | 3 μM |
| Developmental | HGF | 10 ng/mL |
| Developmental | TGFB1 | 10 ng/mL |
| Immune | IFNA | 1.1E4 units/mL |
| Immune | IFNG | 10 ng/mL |
| Immune | IL13 | 1 ng/mL |
| Immune | IL17A | 10 ng/mL |
| Immune | OSM | 10 ng/mL |
| Immune | TNFA | 10 ng/mL |
| Hormonal | Adiponectin | 2 μg/mL |
| Hormonal | Leptin | 10 ng/mL |
| Hormonal | IGF1 | 10 ng/mL |

C

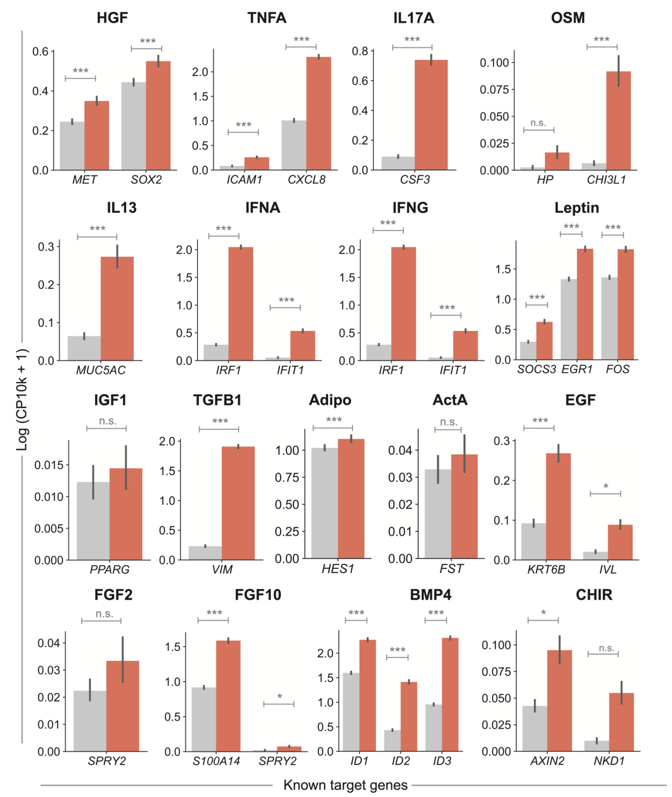


D

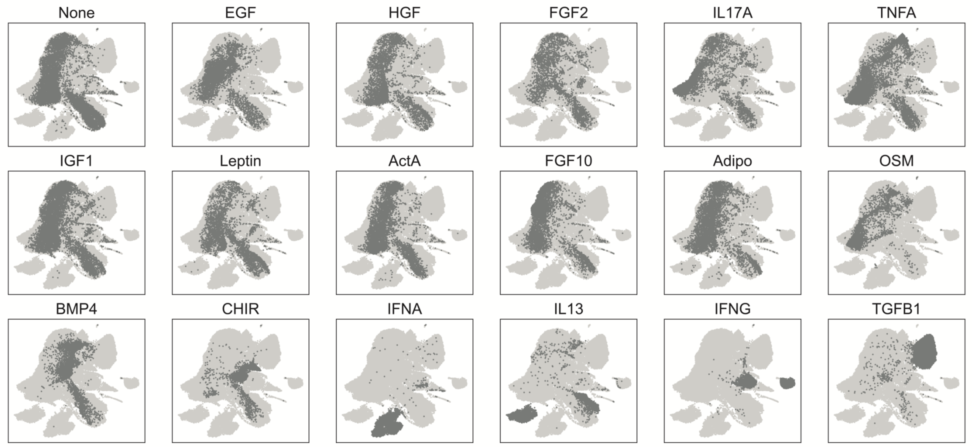


**Supplementary Figure 1. Signaling pathway perturbations in HBEC cultures. Related to Figure 1. (A)** Expression of receptors in HBECs compared to tissue-level expression, colored by selection status (grey: not selected for downstream analysis based on basal cell TPM < 20 and/or expression lower than tissue mean) and manually-assigned receptor signaling category (see methods). Z-score calculated from normalization of TPM in basal cells to mean and standard deviation of expression of gene across human tissues (data from GTex Portal). **(B)** Table of selected ligand and doses used in study. **(C)** Expression of 1-3 previously published target genes in control and signaling perturbation data. CP10k = Counts per 10k total counts. References supporting selection of these target genes is shown in **Table S2**. **(D)** UMAPs of all data shown in 1D, colored individually by each condition in grey with rest of the cells in light grey.

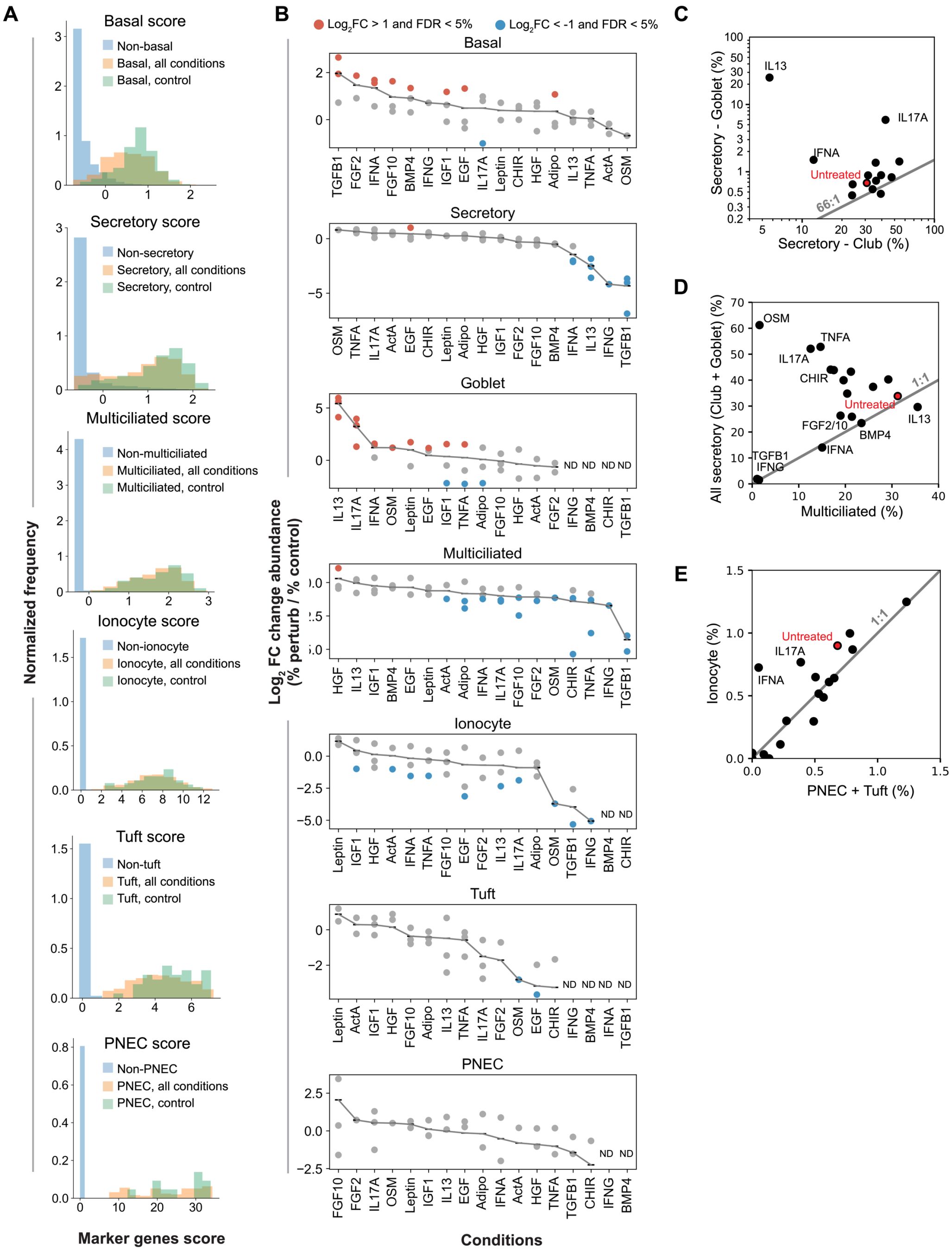


**Supplementary Figure 2. Cell type expression scores and abundances. Related to Figure 2. (A)** As quality-control for cell type classification, an aggregate 'marker gene score' is calculated from the marker genes for each cell type for every cell (see Methods section on 'composite gene scores' and 'cell type classification in treated samples'). The distribution of marker gene scores is shown for cells annotated to the respective cell type in untreated controls (green) and all conditions (orange); and in other cell types (blue) where the score should be significantly lower. **(B)** Fold change abundance (treatment/control) of different cell types in each condition. Each point represents a donor. Red points represent $\log_2$FC > 1 and statistically significant (FDR 5% using Fisher exact test), blue points represent $\log_2$FC < -1 and statistically significant. ND = Not detected. Combined p-values across all donors are given in **Table S3. (C)** Frequency plots of goblet and club cells revealing that these cell types are present in a roughly constant ratio outside of conditions that induce goblet cell hyperplasia. **(D)** Frequency plots of all secretory and multiciliated cells reveal that goblet cell hyperplasia is associated with variable, signaling-specific changes in secretory:multiciliated cell ratio. **(E)** Frequency plots of ionocytes and PNEC+Tuft cells across all conditions indicate a conserved ratio of rare cell types.

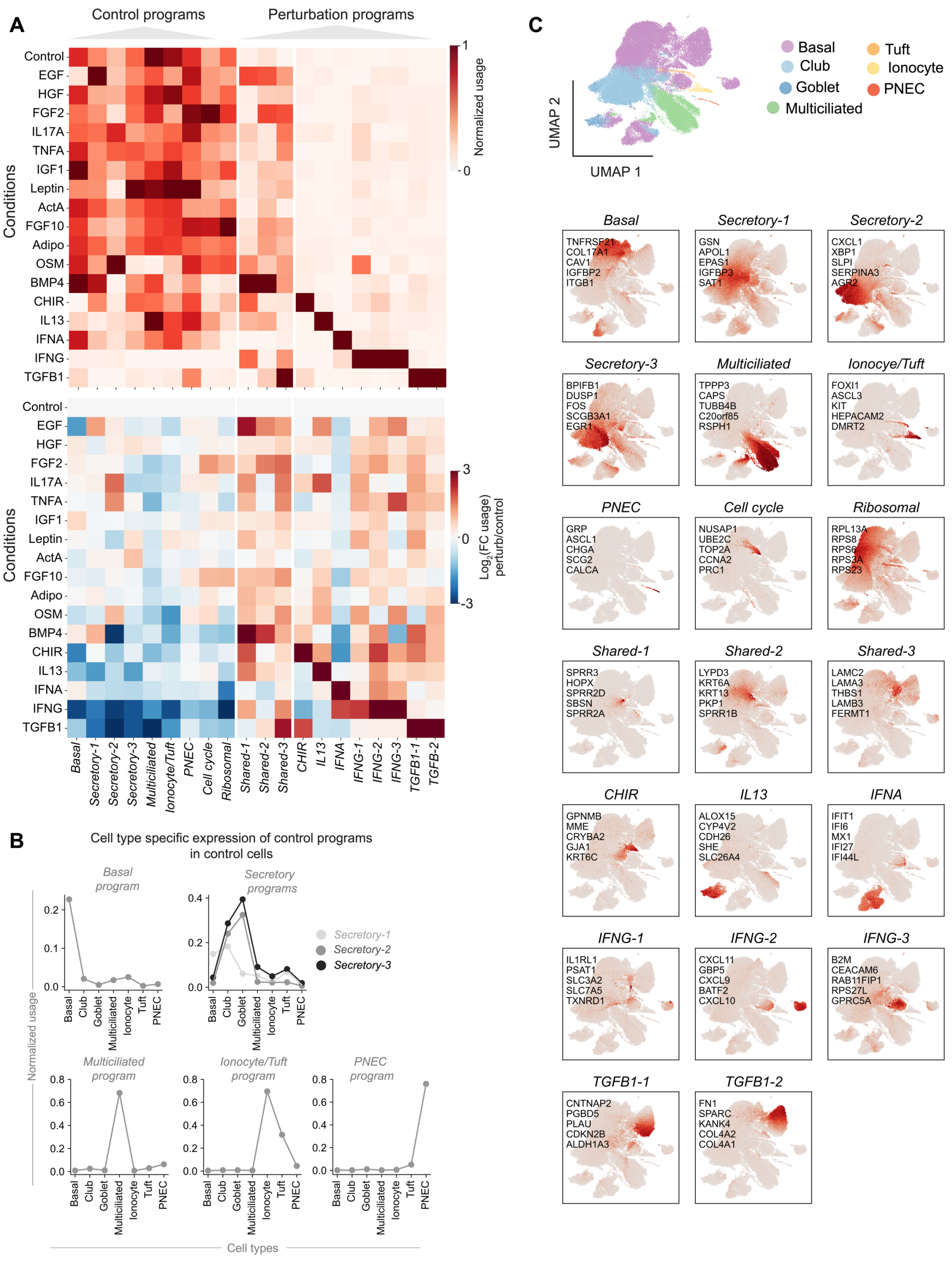


**Supplementary Figure 3. Expression of cNMF programs across samples. Related to Figure 3. (A)** Top: Normalized usage of all gene programs, extending **Fig. 3C** to include programs present in the untreated controls. See **Fig. 3C** legend for heatmap normalization. Bottom: $Log_2$ fold change in usages comparing treatment with untreated samples. **(B)** Cell type specific usage of control programs in untreated cells, extending **Fig. 3D** to programs present in the untreated controls. **(C)** Normalized usage of all gene programs plotted on a UMAP of all cells.

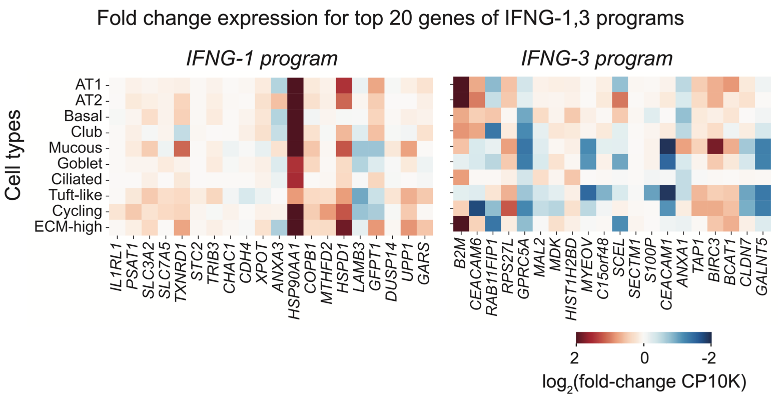


**Supplementary Figure 4. Enrichment of IFNG response genes in cell types from severe COVID-19 patient samples. Related to Figure 5.** Heatmap shows fold change in expression of top 20 genes of *IFNG-1* and *IFNG-3* programs in Covid disease versus control samples. To ensure that no single gene dominates the enrichment score shown in Fig. 5, the enrichment scores represent the median score of 100 bootstraps of 7 of the 20 genes shown here (see Methods for details).

#### Supplementary Tables

##### Table S1. Previous reported responses of the mucociliary airway epithelium to the signaling pathway responses investigated in this paper

| Stimulus | Published Effect in Lung Culture Systems | Selected References |
|---|---|---|
| **EGF** | Squamous metaplasia and epithelial-to-mesenchymal transition(EMT) | 17,63 |
| **HGF** | Promoted cell motility during wound repair and increased differentiation | 64,65 |
| **FGF2** | Decreased proliferation and control of FGF signaling in the trachea | 66–68 |
| **IL17A** | Inflammation and goblet cell metaplasia | 12,20,69 |
| **Adiponectin** | Increased proliferation and wound healing response | 70–72 |
| **TNFA** | Inflammation including goblet cell metaplasia, EMT | 73,74 |
| **IGF1** | Unknown; knockout studies suggest role in inflammation and epithelial maturation | 75 |
| **Leptin** | No effect observed in immortalized hBECs | 70 |
| **Activin A** | Induced in hBECs by cigarette smoke but no evidence for direct effect of signaling ligand. Member of TGF-b family, so may impact similar phenotypes. | 76 |
| **FGF10** | Mediates epithelial cell proliferation in adult homeostasis and repair after injury, can result in increased basal cell colony formation and proliferation | 67,68,77 |
| **BMP4** | Suppresses basal cell proliferation and differentiation, induces KRT6A/B, IVL, KRT14, SFN in ALI cultures, leading to squamous metaplasia | 43,78–80 |
| **CHIR99021 (GSK3 inhibitor)** | Modulates canonical Wnt signaling along with other effects. Prevents normal differentiation and drives a hyperproliferative state. In COPD-derived ALI-HBECs, Wnt inhibited epithelial differentiation, polarity and barrier function, and induced TGF-β-related epithelial-to-mesenchymal transition (EMT). | 81–84 |
| **IFNA** | Type I interferon. hBEC response to treatment with IFNA leads to expansion of goblet cells (MUC5B expression) at the expense of multiciliated cells. However, treatment of murine airway cells with IFNA led to a decrease in secretory cells. | 85 |
| **IFNG** | Type II interferon, involved in viral host defense. Most studies looking at hBEC response focus on chemokine secretion. Interferon gamma signaling is upregulated in vivo in response to COVID infection and associated with loss of normal epithelial differentiation and accumulation of metaplastic basal cells | 86,87 |
| **IL13** | Induction of mucus metaplasia and mucus hypersecretion | 19,88 |
| **OSM** | Loss of barrier function, mucus metaplasia, and activation of inflammatory pathways via STAT activation in human airway cells. | 36,89 |
| **TGFB1** | Cell cycle arrest, p53 signaling, cell spreading, squamous metaplasia, and EMT | 90–92 |

**Table S2. References for gene targets of signaling molecules used in the study (supporting Fig. S1C)**

| Stimulus | Target Genes | References |
|---|---|---|
| EGF | *KRT6B, IVL* | [17] |
| HGF | *MET, SOX2* | [93] |
| FGF2 | *SPRY2* | [94] |
| IL17A | *CSF3* | [95] |
| Adiponectin | *HES1* | [96] |
| TNFA | *ICAM1, CXCL8* | [97] |
| IGF1 | *PPARG* | [98] |
| Leptin | *SOCS3, EGR1, FOS* | [99] |
| Activin A | *FST* | [76] |
| FGF10 | *S100A13, S100A14, SPRY2* | [100] |
| BMP4 | *ID1, ID2, ID3* | [101] |
| CHIR99021 (GSK3 inhibitor) | *AXIN2, NKD1* | [102] |
| IFNA | *IFIT1, IRF7* | [103,104] |
| IFNG | *IFIT1, IRF1* | [104,105] |
| IL13 | *MUC5AC* | [19] |
| OSM | *HP, CHI3L1* | [36] |
| TGFB1 | *VIM* | [92] |

**Table S3. Frequencies of cell types (supporting Fig. 2B and S2B)**. See attached Excel spreadsheet.
Worksheets:
Sheet S3-1 (Fold changes): $Log_2$ fold change of relative abundance of cell types between signaling condition and untreated sample averaged between donors. Donor specific p-value was calculated using Fisher's Exact test and integrated by Fisher's method. FDR calculated using Benjamini Hochberg method.
S3-2 (Frequencies donor 1): Frequencies of cells in all annotated cell type for donor 1
S3-3 (Frequencies donor 2): Frequencies of cells in all annotated cell type for donor 2
S3-4 (Frequencies donor 3): Frequencies of cells in all annotated cell type for donor 3

**Table S4. Differentially expressed genes upon stimulation of 17 signaling pathways, analyzed separately for each canonical cell type (supporting Fig. 3A)**. See attached Excel spreadsheet. These include number of genes showing >2-fold differential expression and FDR < 0.05 (rank sum test, Benjamini Hochberg correction) in basal, secretory (club, goblet), multiciliated and rare (ionocyte, tuft, PNEC) cells following each treatment.

**Table S5. Usages of cNMF programs for in our dataset (supporting Fig. 3)**.
See attached Excel spreadsheet. Worksheets:
S5-1 (Averaged usage): Usage averaged per condition and annotated cell type.
S5-2 (Usage per cell): Usage per cell for all cells in the data

**Table S6. Gene loadings for cNMF programs for 3000 highly variable genes (supporting Fig. 3)**.
See attached Excel spreadsheet. Worksheets:
S6-1 (Top 20 genes): Top 20 genes for each program
S6-2 (All gene loadings): Gene loadings for all 3000 genes
